# Image processing tools for petabyte-scale light sheet microscopy data

**DOI:** 10.1101/2023.12.31.573734

**Authors:** Xiongtao Ruan, Matthew Mueller, Gaoxiang Liu, Frederik Görlitz, Tian-Ming Fu, Daniel E. Milkie, Joshua L. Lillvis, Alexander Kuhn, Johnny Gan Chong, Jason Li Hong, Chu Yi Aaron Herr, Wilmene Hercule, Marc Nienhaus, Alison N. Killilea, Eric Betzig, Srigokul Upadhyayula

**Author notes:** These authors contributed equally, names are listed alphabetically (Eric Betzig, Srigokul Upadhyayula).

## Abstract

Light sheet microscopy is a powerful technique for high-speed 3D imaging of subcellular dynamics and large biological specimens. However, it often generates datasets ranging from hundreds of gigabytes to petabytes in size for a single experiment. Conventional computational tools process such images far slower than the time to acquire them and often fail outright due to memory limitations. To address these challenges, we present PetaKit5D, a scalable software solution for efficient petabyte-scale light sheet image processing. This software incorporates a suite of commonly used processing tools that are memory and performance-optimized. Notable advancements include rapid image readers and writers, fast and memory-efficient geometric transformations, high-performance Richardson-Lucy deconvolution, and scalable Zarr-based stitching. These features outperform state-of-the-art methods by over one order of magnitude, enabling the processing of petabyte-scale image data at the full teravoxel rates of modern imaging cameras. The software opens new avenues for biological discoveries through large-scale imaging experiments.

## 1 Introduction

Light sheet microscopy enables fast 3D imaging of cells, tissues, and organs [1]. Within this realm, variants like multi-view selective plane illumination microscopy [2–4], lattice light sheet microscopy (LLSM) [5], axially swept light sheet microscopy [6, 7], and single objective light sheet microscopy [8–10], offer higher resolution and fast imaging speed. Combined with expansion microscopy [11], these techniques have been used to millimeter-scale or larger cleared and expanded specimens [12, 13], while achieving nanoscale resolution. In such cases, the data produced from a single experiment can explode to the petabyte range. These high data generation rates introduce significant challenges for data storage and processing that complicate visualization, assessment, and analysis. First, even individual volumes from a 4D time series can be so large as to render their pre-processing unwieldy or impossible for conventional processing codes. Second, acquisition in a non-Cartesian coordinate space adds substantial computational overhead. Third, light sheet data is often acquired at multi-terabyte-per-hour rates which is too fast for conventional tools to process in real time. This impedes the rapid feedback needed to adjust imaging conditions or locations on the fly or to extract biological insights from the resulting datasets in a timely manner.

Numerous computational tools encompassing various functionalities have been developed to facilitate light sheet image pre-processing, including deskew and rotation [14, 15], deconvolution [16, 17], stitching [18, 19], and visualization [20, 21]. While these tools have proven valuable for light sheet images on the gigabyte scale, their utility wanes for data sizes surpassing the terabyte threshold, due to a lack of scalability and efficiency required to process images in real-time. Furthermore, many of these tools are standalone applications, providing only partial processing steps in a specialized context and varying input formats and requirements. This situation often requires extensive manual effort to integrate them into multi-step workflows, limiting their utility, especially for large-scale data.

To address these challenges, particularly for long-term imaging of subcellular dynamics or vast multicellular image volumes, we developed PetaKit5D, a software solution designed to enable real-time processing of petabyte-scale imaging data. The software contains commonly used pre- and post-processing tools that are memory and performance-optimized, including deskew, rotation, deconvolution, and stitching, all integrated into a high-performance computing framework capable of executing user-defined functions in a scalable and distributed manner.

To further increase throughput, we developed novel algorithms for image input/output (IO) using the Zarr data format for image storage [22] and processing in conjunction with custom parallelized image readers and writers. These capabilities have been optimized for partitioned parallel processing of petabyte-scale datasets. The software incorporates an online mode during image acquisition to automatically process data and provide near-instantaneous feedback that is critical during long-term time series or high-throughput large sample imaging.

We developed PetaKit5D in MATLAB and offer Python wrappers for the deployed version. To ensure accessibility for users with little or no programming experience, the software includes a user-friendly graphical user interface (GUI).

## 2 Results

### 2.1 Overall design: distributed computing accelerates image processing

High frame rate modern cameras enable light sheet microscopes to capture images at nearly four terabytes per hour per camera. This presents formidable challenges for sustained image acquisition, real-time (de)compression, storage, and processing, especially when using a single conventional workstation. In response, we developed a distributed computing architecture comprising a cluster of computing nodes and networked data storage servers that enables uninterrupted streaming and real-time processing of vast quantities of data continuously acquired over extended periods. Our standard workflow is illustrated in Fig. 1a.

**Fig. 1:**
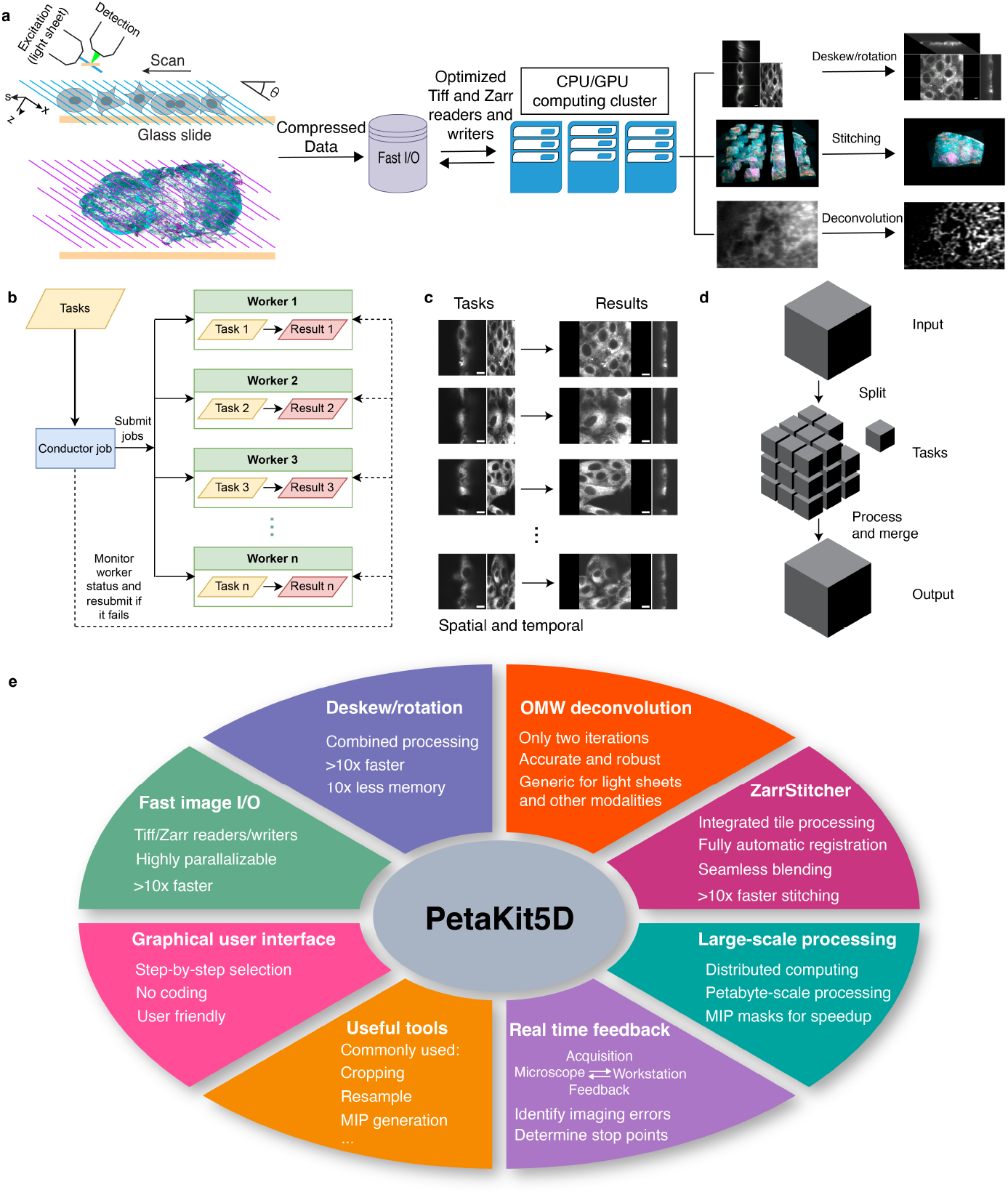
Overall design of the image processing framework. **a**, image acquisition and processing workflows. **b**, illustration of the generic distributed computing framework. **c**, illustration of distributed processing of many independent files across multiple workers. **d**, illustration of distributed processing of the splitprocess-merge mechanism for the distributed processing of a large image file. **e**, overall functionalities and features in PetaKit5D. OMW is an acronym for OTF Masked Wiener Richardson-Lucy deconvolution method.

Our approach uses a generic distributed computing framework in MATLAB to parallelize user-defined functions ((Fig. 1b). The complete dataset or set of tasks is divided into distinct, self-contained subtasks, each appropriately sized for processing by an individual worker unit with one or more CPU cores or GPUs (Fig. 1c-d). A conductor job orchestrates all operations, distributes tasks across the computing cluster, and monitors their progress to completion (Fig. 1b). Failed jobs are automatically resubmitted with additional resources. Our MATLAB-based framework offers greater flexibility for various task types, enhanced robustness against failures, and seamless integration across multiple processing steps, compared with Spark [23] and Dask [24]. We use it to manage all processing methods in PetaKit5D (Fig. 1e).

### 2.2 Fast image readers and writers

Efficient image reading and writing are essential for real-time image processing. The widely used Tiff format stores 2D and 3D microscopy data, offers the ability to compress (such as LZW (Lempel-Ziv-Welch) compression), and can include specialized metadata (such as OME-TIFF (Open Microscopy Environment TIFF) [25]). Unfortunately, conventional image readers and writers for the Tiff format are not designed for large-scale compressed data, being restricted to single-threaded operations (e.g., libtiff). For instance, an 86 GiB 16-bit Tiff file (512 × 1, 800 × 50,000) with libtiff (LZW compression) takes approximately 8.5 and 16 minutes to read and write, respectively (Fig. S1a-b, last groups). These speeds pose a considerable bottleneck for efficient image processing, especially when the entire image needs to be loaded for the processing. Memory mapping is an alternative technique to process large images (i.e., tifffile [26] in Python), but it is mostly limited to working with uncompressed data in their native byte order. This limitation can significantly increase storage requirements for large datasets, and the processing may still be constrained by slow read and write speeds.

To rectify this, we developed an optimized Tiff reader and writer in C++. This implementation leverages the OpenMP framework [27] to facilitate concurrent multi-threaded reading and writing (where only the compression process is parallelized in writing). Our Cpp-Tiff reader and writer are over 22× and 7× faster than conventional ones, respectively, for compressed data (Fig. 2a-b and Fig. S1a-b for a 24-core node). Moreover, they substantially outperform the fast Python reader and writer library for Tiff files (tifffile [26] in Python, Fig. 2a-b, and Fig. S1 a-b). Their speeds also increase linearly as more CPU cores are devoted to read/write operations (Fig. S1e-f).

**Fig. 2:**
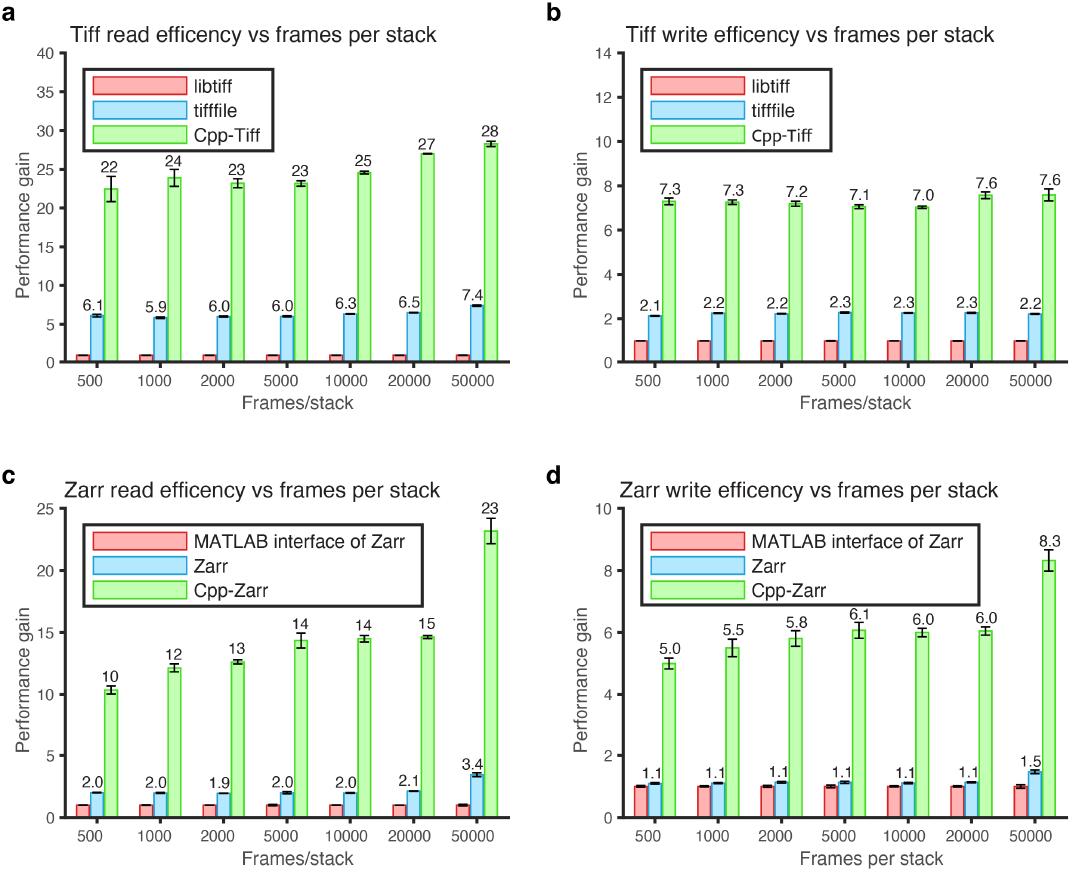
Performance improvement factors of our C++ Tiff and Zarr readers and writers. **a**, performance gains of our Cpp-Tiff reader versus the conventional Tiff reader in MATLAB and tifffile reader in Python versus the number of frames in the 3D stack. **b**, performance gains of Cpp-Tiff writer versus the conventional tiff writer in MATLAB and the tifffile writer in Python versus the number of frames in the 3D stack. **a**, performance gains of our Cpp-Zarr reader versus the conventional Zarr reader (MATLAB interface of Zarr) and native Zarr in Python versus the number of frames in the 3D stack. **d**, performance gains of Cpp-Zarr writers versus the conventional Zarr writer (MATLAB interface of Zarr) and native Zarr in Python versus the number of frames in the 3D stack. The image frame size is 512 × 1800 (xy) in all cases. The benchmarks were run independently ten times on a 24-core CPU computing node (dual Intel Xeon Gold 6146 CPUs).

Although the Tiff format is commonly used for raw microscopy images, it is not the most efficient for parallel reading and writing, especially for very large image datasets. One major limitation is its single-container structure for file writes, which restricts it to single-threaded operations. To overcome this, we instead chose Zarr [22], a next-generation file format optimized for multi-dimensional data. Zarr efficiently stores data in non-overlapping chunks of uniform size (border regions may be padded to match the full chunk size) and saves them as individual files. The format is similar to N5 [28], OME-Zarr [29], and TensorStore [30].

Zarr allows individual jobs to access only the specific region of interest at a given time. Distinct regions can be saved to disk independently and in parallel. Using optimized C/C++ code that leverages OpenMP, our Zarr reader/writer is 10-23× faster for reading and 5-8× faster for writing (Fig. 2c-d, Fig. S1c-d) than the conventional implementation (using MATLAB’s *blockedImage* function to interface with the Python version of Zarr). Their performances also scale as more CPU cores are devoted to read/write operations (Fig. S1g-h). Our implementation is also 5-10× and 5-8× faster for read/write compared with the native Python implementation of Zarr (Fig. 2c-d, Fig. S1c-d). Moreover, compared to TensorStore, Cpp-Zarr is 2.2× and 1.5× faster for reading and writing, respectively, for their preferred data orders (row-major in TensorStore and column-major in Cpp-Zarr, Table S1). We opted to use the zstd compression algorithm [31] at compression level 1 to achieve better compression ratios at comparable read/write times to the lz4 algorithm [32] at level 5 (default in native and OME-Zarr) (Fig. S1i-k). We also created a Parallel Fiji Visualizer plugin that quickly opens compressed Tiff and Zarr files using our fast readers, enabling efficient data visualization and inspection in Fiji [33] (Supplementary note 3).

### 2.3 Fast combined deskew and rotation

In many light sheet microscopes, the excitation and detection objectives are oriented at an angle with respect to the substrate holding the specimen. It is convenient in such cases to image the specimen by sweeping it in the plane of this substrate, but the resulting raw image stack is then sheared and rotated with respect to the conventional specimen Cartesian coordinates (Fig. 3a). Traditionally, the data is transformed back to these coordinates by deskewing and rotating in two sequential steps (Fig. 3b). However, zero padding during deskew drastically increases data size, slowing computation and risking out-of-memory faults, particularly for large images with many frames (Fig. 3c).

**Fig. 3:**
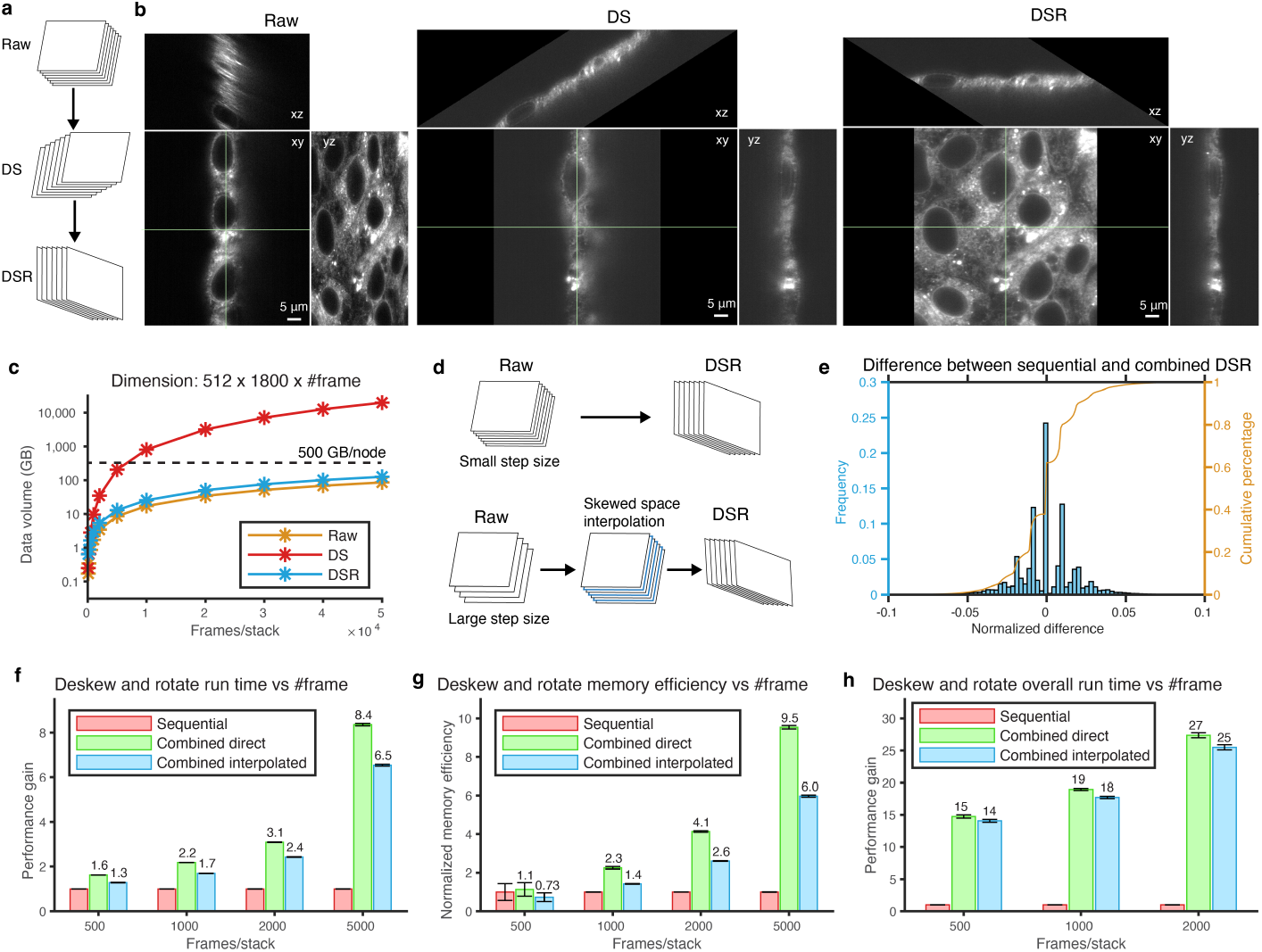
Combined deskew and rotation. **a**, traditional sequential deskew and rotation processes. **b**, orthogonal views of raw (skewed), deskewed (DS), and deskewed/rotated (DSR) images for cultured cells. **c**, the semi-log plot of the data sizes of a stack of raw 512 × 1800 pixel images, deskewed, deskewed/rotated data as a function of the number of frames per stack. The dashed line indicates the memory limit (500 GB) of a computing node. **d**, the addition of skewed space interpolation prior to deskew/rotation when the z-step size between image planes is too large. **e**, histogram of the normalized differences between deskewed/rotated results of sequential versus combined methods for the same image in **b. f**, performance gain for combined direct and combined interpolated over sequential deskew/rotation versus the number of frames per stack, from 500 to 5,000. Comparisons do not include read/write time, which is considered in panel **h. g**, memory efficiency gain for the same three scenarios. **h**, performance gain for the same three scenarios. This comparison does include the differences in read/write time when the conventional Tiff software is used for the sequential deskew/rotation, and our Cpp-Tiff is used for combined deskew/rotation.

To address this issue, we combined deskew and rotation into a single step, which is possible given that both operations are rigid geometric transformations. While prior studies have explored combining processing techniques using vertical interpolation in the deskew/rotated space and customized transformations [9, 10, 34], they are limited either in speed or the amount of data they can handle. Unlike previous approaches, our method first interpolates the data in the raw skewed space (depending on the scan step size), followed by standard affine transformation. When the ratio of the scan step size in the xy plane to the xy voxel size (defined as “skew factor”) is smaller than 2.0, this is readily feasible (Fig. 3d, top). However, when the skew factor is larger than 2.0, artifacts may manifest due to the interpolation of voxels that are spatially distant within the actual sample space during the combined operations, as depicted in Fig. S2a-b. Thus, in this case, we first interpolate the raw skewed data between adjacent planes within the proper coordinate system to add additional planes to reduce potential artifacts in the following combined operations (Fig. 3d, bottom).

Our combined deskew and rotation method yield nearly identical results to the same operations performed sequentially (Fig. 3e and Fig. S2c-e). The combined operation becomes increasingly fast and memory efficient compared to sequential operation as the number of frames increases. This enables us to process 10× larger data with the same computational resources (Fig. 3f-g, and Fig. S2f-g). By additionally combining our fast Tiff reader/writer with combined deskew/ rotation, we achieve a 20× gain in processing speed compared to conventional Tiff and sequential processing, allowing us to process much larger data (Fig. 3h and Fig. S2h). Our approach is significantly faster than the CPU implementation in pyclesperanto [35]) and can handle larger volumes containing many frames (Fig. S2i). Finally, resampling and cropping, if necessary, can also be integrated with deskewing and rotation to optimize processing efficiency and minimize storage requirements for intermediate data.

### 2.4 OTF masked Richardson-Lucy deconvolution

Deconvolution plays a crucial role in reconstructing the most accurate possible representation of the sample from light microscopy images, especially for light sheet images with strong side lobes associated with higher axial resolution [36]. Richardson-Lucy (RL) deconvolution is the most widely used approach due to its accuracy and robustness [37, 38]. We have found that applying RL to the raw light sheet data prior to combined deskew/rotation is not only faster (due to no zero padding) but also yields better results with fewer edge artifacts (Fig. S3a). To do so, the reference point spread function (PSF) used for deconvolution must either be measured in the skewed space as well (Fig. S3c), or else calculated by skewing a PSF acquired in the sample Cartesian coordinates (Fig. S3b).

RL is an iterative method that typically requires 10-200 iterations (Biggs accelerated version [39]) to converge, depending on the type of light sheet or image modalities. Consequently, RL deconvolution is the most computationally intensive step for large datasets relative to deskew, rotation, and stitching, even with GPU acceleration. Despite being notably faster than the traditional RL method, the Wiener-Butterworth (WB) method proposed by Guo et al. [17] with unmatched backward projector was initially demonstrated on Gaussian light sheets with ellipsoid support, and failed to achieve full-resolution reconstruction of LLSM images (Fig. 4a-b, Fig. S3d-f) since it truncates the Optical Transfer Function (OTF) near the edges of its support (Fig. 4a-b, Fig. S3d-f), resulting in the loss of information. This limitation is particularly pronounced for lattice light sheets capable of high axial resolution, such as the hexagonal, hexrect, and multi-Bessel types [36], whose OTF supports are nearly rectangular rather than ellipsoidal in the xz and yz planes. Another concern is that the WB method suppresses high-frequency regions near the border of its back projector’s ellipsoid, thereby underweighting or even eliminating high-resolution information in the deconvolved images.

**Fig. 4:**
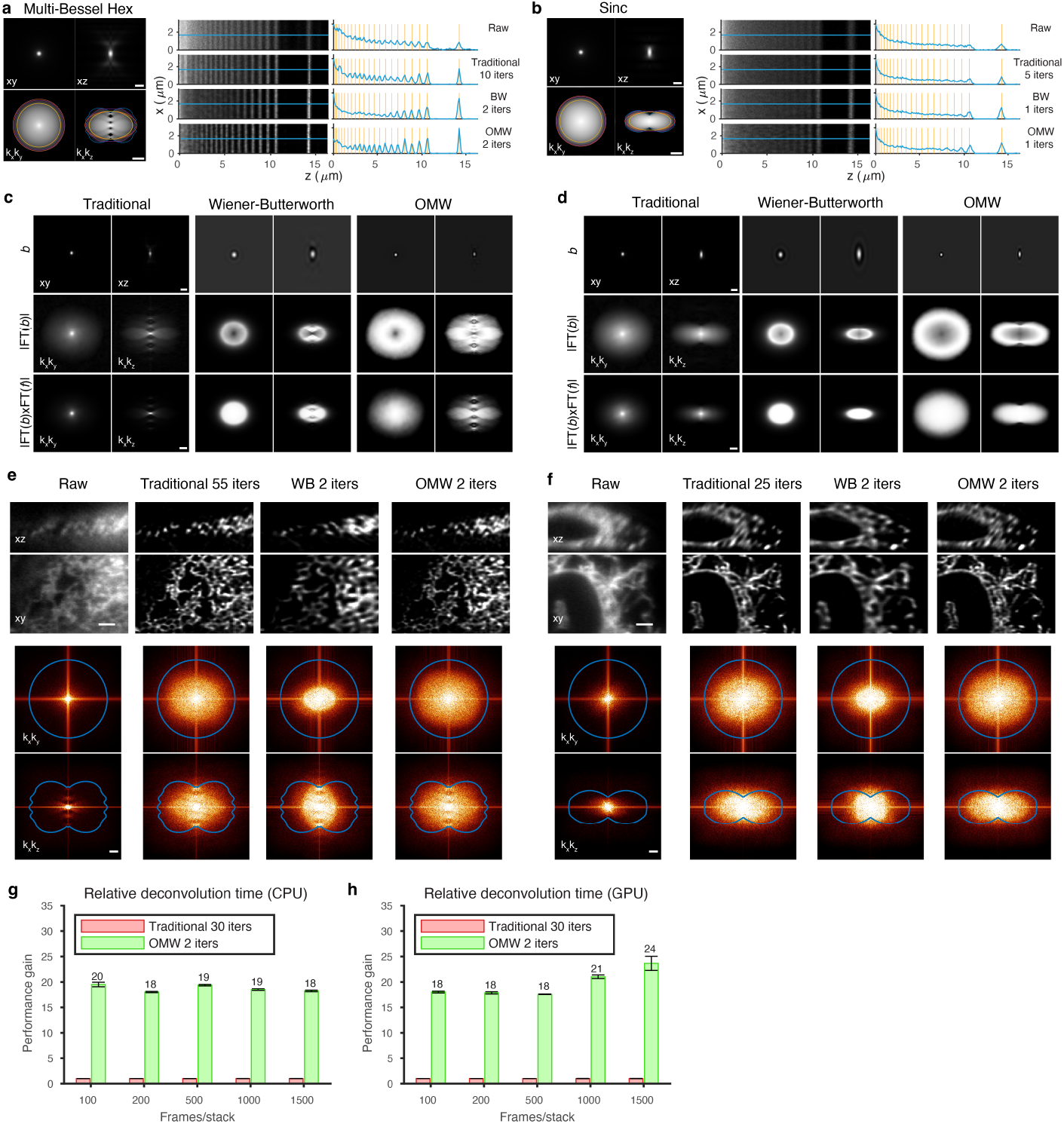
Fast RL deconvolution. **a**, Left: theoretical xy and xz PSFs (top, intensity *γ* = 0.5, scale bar: 1 *µm*) and OTFs (bottom, log-scale, scale bar: 2 *µm*^−1^) for the multi-Bessel (MB) light sheet with excitation NA 0.43 and annulus NA 0.40 - 0.47. Blue: theoretical support; orange and yellow: theoretical maximum (orange) and experimental (yellow) envelopes for the WB method; magenta: experimental envelope for the OMW method. Right: illustration of deconvolution of a simulated stripe pattern. The raw and deconvolved images with traditional, WB, and OMW methods are displayed along with their line cuts. The orange lines indicate the theoretical line locations, and the blue curves give the actual intensities along the line cuts. **b**, similar results for a Sinc light sheet (NA 0.32, *σ*_*NA*_ = 5.0). **c-d**, illustration of backward projectors (top, scale bar: 1 *µm*), their Fourier spectra (middle, intensity *γ* = 0.5), and the products with forward projectors in Fourier spaces (bottom, intensity *γ* = 0.5, scale bar: 1 *µm*^−1^) for the MB light sheet (**c**) and Sinc light sheet (**d**). **e-f**, orthogonal views of cell images for raw, traditional RL, WB, and OMW methods for the MB light sheet (**e**) and Sinc light sheet (**f**), with iteration numbers as shown (scale bar: 2 *µm*). The Fourier spectra are shown below each deconvolved image (intensity *γ* = 0.5, scale bar: 1 *µm*^−1^). **g-h**, relative deconvolution acceleration for traditional RL and OMW methods on CPU (**g**) and GPU (**h**) (only the deconvolution performance is shown in the comparison). Each test in **g** and **h** was run independently ten times.

To address these issues, inspired by Zeng et al. [40] and Guo et al. [17], we optimized the backward projector by using the convex hull of the OTF support to define an apodization function. This function filters noise close to the support and eliminates all information beyond it (Fig. 4a-b, Fig. S3d-f, Fig. S4a-b), and is applied to the Wiener filter (Fig. S4c-d). Unlike the WB method, this OTF masked Wiener (OMW) technique covers all relevant frequencies in the Fourier space (Fig. 4c-d, Fig. S3g-h) and achieves full-resolution image reconstruction while maintaining rapid convergence speed (Fig. 4e-f, Fig. S3i-l). By using the OTF support for apodization, the OMW method is generic for any PSF. Our specific implementation offers a 10-fold speed improvement compared to the traditional RL method (Biggs version) on both CPUs and GPUs (Fig. 4g-h, Fig. S5a-f). A detailed comparison with other deconvolution methods is provided in Supplementary Note 4.

The performance of RL relies crucially on finding an optimum number of iterations: too few yields fuzzy images and, in the case of LLSM, incomplete sidelobe collapse; too many amplifies noise and potentially collapses and fragments continuous structures as represented in the deconvolved image. To find an optimum, we use Fourier Shell Correlation (FSC) [41]. While the traditional RL method often needs tens of iterations to optimize resolution by the FSC metric (Fig. S5g), the OMW method typically only needs two iterations when we use FSC to determine the optimal Wiener parameter (Fig. S5h-i) for the backward projector for light sheet images. Widefield and confocal images, however, may need more iterations (Fig. S6).

### 2.5 ZarrStitcher: Zarr-based scalable stitching

To image specimens such as organoids, tissues, or whole organisms larger than the field of view of the microscope, it is necessary to stitch together multiple smaller image tiles. Overlap regions between adjacent tiles facilitate precise registration and stitching. With the combination of high-resolution light sheet and expansion microscopy, thousands of tiles comprising hundreds of terabytes of data may be generated. This presents substantial challenges for existing stitching software, particularly with respect to the large number of tiles, the overall data size, and the need for computational efficiency. To address these issues, we developed ZarrStitcher, a petabyte-scale framework for image stitching.

ZarrStitcher involves three primary steps (Fig. 5a): data format conversion, cross-correlation registration, and stitching (fusion). We first convert tiles into the computationally efficient Zarr format, while also applying user-defined preprocessing functions such as flat-field correction and data cropping if necessary. Next, we use the normalized cross-correlation algorithm [42] to correct for sample movement and stage motion errors and thereby accurately register the relative positions of adjacent tiles. We then apply a global optimization to infer the optimal shifts (Fig. 5b, middle) of all tiles collectively. This better manages potential discrepancies between neighboring tiles than the typical “greedy” local approach (Fig. 5b, left). For large volumes, data is often collected in multiple batches, each consisting of multiple tiles, sometimes with differing rectangular grids in each batch. In such cases, we implement two-step optimization, where global optimization is first applied to each batch, followed by optimization across batches (Fig. 5b, right).

**Fig. 5:**
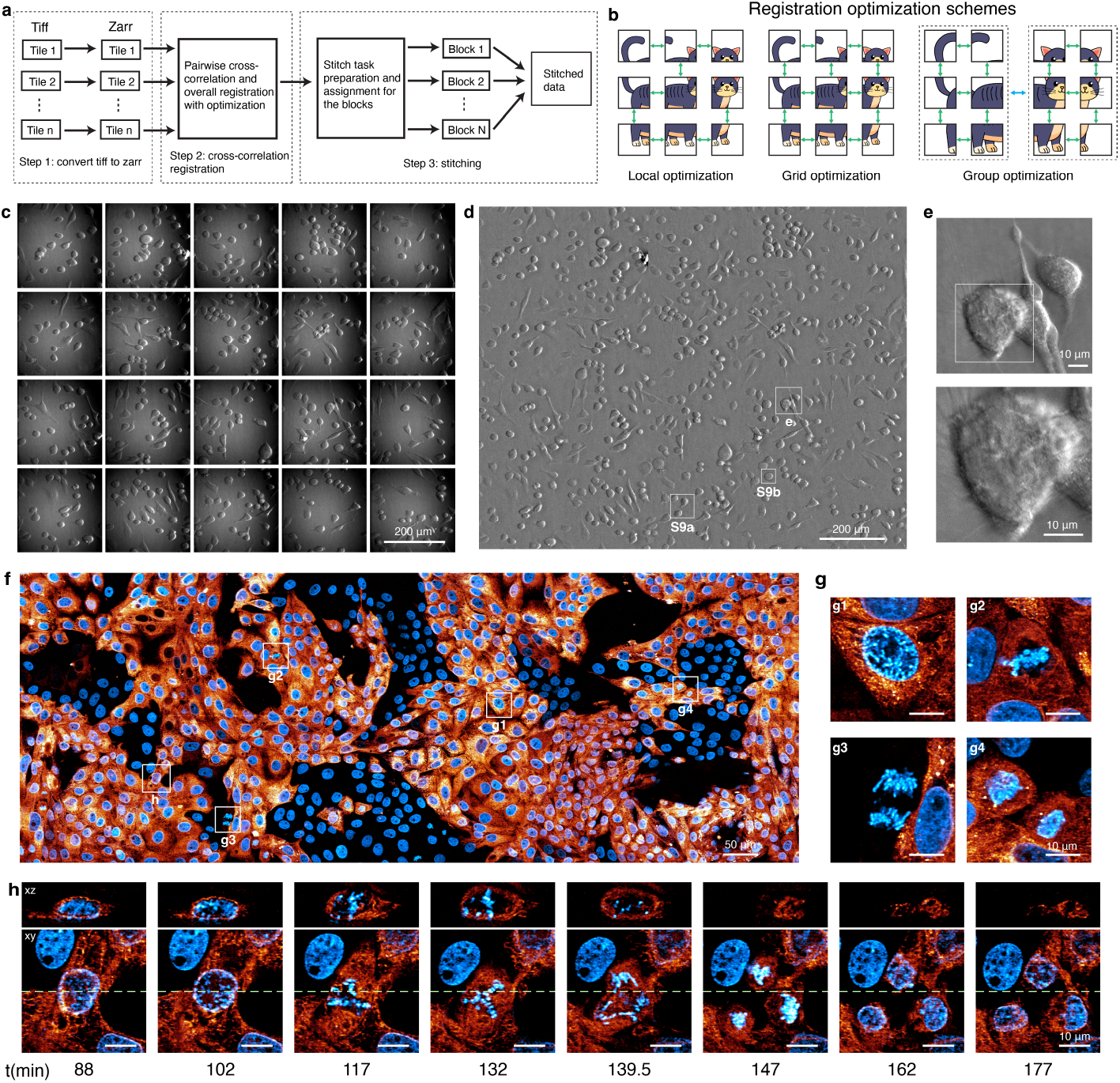
The Zarr-based distributed stitching framework. **a**, schematic of the stitching steps. **b**, schematic of different registration methods: local, grid, and grouped. Green or blue double arrow lines indicate pairs of tiles/groups involved in registration. **c**, raw 2D oblique illumination “phase” tiles of live HeLa cells before processing. **d**, final processed phase image after flat-field correction, stitching, and deconvolution. Boxes labeled S9a and S9b indicate regions shown at higher magnification in Fig. S9a and S9b. **e**, zoomed region of **d** showing retraction fibers. **f**, xy MIP view of long-term large field of view 3D imaging of cultured LLC-PK1 cells. Blue: H2B-Cherry; hot: Connexin-Emerald. Intensity *γ* = 0.5. **g**, cropped regions from **e** showing stages in cell division. **h**, time-lapse orthogonal views of one cell division into three daughter cells from **f**. The green dashed lines indicate the orthogonal slice positions in the xz plane.

The final operation in ZarrStitcher involves stitching the registered tiles together into a single unified volume. We developed a scalable distributed architecture to this end, with individual tasks allocated to different workers for different subregions. The software incorporates multiple methods to address overlapping regions, including direct merging, mean, median, or feather blending. Feather blending, a type of weighted averaging with weights determined by distances to the border, has proven to be particularly effective [43].

ZarrStitcher is substantially faster than BigStitcher-Spark (Spark version of BigStitcher) [19] (Table S2): in the case of the 108 TiB dataset for the entire mouse brain imaged with 4× expansion using ExA-SPIM ([13]), ZarrStitcher took 1.4 hours using 20 computing nodes (480 CPU cores) to assemble the complete volume, 14.3× faster than BigStitcher-Spark. This is an active research area, with ongoing development efforts working to close the performance gap (Table S2). Stitching-spark [12], another alternative, is not usable at this scale, due to its use of Tiff files that are limited to 4 GB size. ZarrStitcher out-performs BigStitcher-Spark in fusing images in cases with extensive overlap, minimizing ghost image artifacts caused by imperfect structure matches in overlapping regions (Fig. S7).

By integrating fast readers and writers, combined deskew and rotation, and ZarrStitcher, we assembled a pipeline with real-time feedback during microscopy acquisition that facilitates rapid analysis and decision-making. In the online processing mode, this pipeline uses the native coordinates for stitching without global registration. It allows acquisition errors to be identified mid-stream, so that corrections can be made (Fig. S8a) and helps determine when the specimen has been fully imaged so the acquisition can be concluded (Fig. S8b). It also enables quick identification of specific cells or specific events in a large field view worthy of more detailed investigation (Fig. 5c-e), such as cell fusion (Fig. S9a) or cell division (Fig. S9b, Movie S1).

We have also coupled our processing pipeline to NVIDIA’s multi-GPU IndeX platform [27] to enable real-time visualization of 4D petabyte-scale data at full resolution (Supplementary Note 5, Movie S3). This allows us to simultaneously follow the dynamics of hundreds to thousands of cells (Fig. 5f, Movies S2 and S3), and identify infrequent or rare events such as normal cell divisions or the division of a cell into three daughter cells (Fig. 5g-h, Movies S2 and S3). Furthermore, it enables us to explore their 3D high-resolution subcellular structures in detail over an extended period (Movies S2 and S3). The entire processing and imaging pipeline is applicable to many microscope modalities in addition to light sheet microscopy. These include high speed, large field of view oblique illumination “phase” imaging (Fig. 5c-e, Movie S1), large volume adaptive optical two-photon microscopy (Fig. S9c-f, Movie S4), widefield imaging (Fig. S6a-b), and confocal imaging (Fig. S6c-d).

### 2.6 Strategies for large-scale processing

For large datasets consisting of many tiles, it is most efficient to stitch the tiles in skewed space before deconvolution (Fig. 6a), thereby eliminating duplicated effort in overlap regions as well as potential edge artifacts. Deconvolving still in skewed coordinates immediately thereafter is most efficient (Fig. S3a), because the data is more compact than after deskewing. Thus, the optimal processing sequence is stitching (if necessary), followed by deconvolution, and finally combined deskew and rotation (Fig. 6a).

**Fig. 6:**
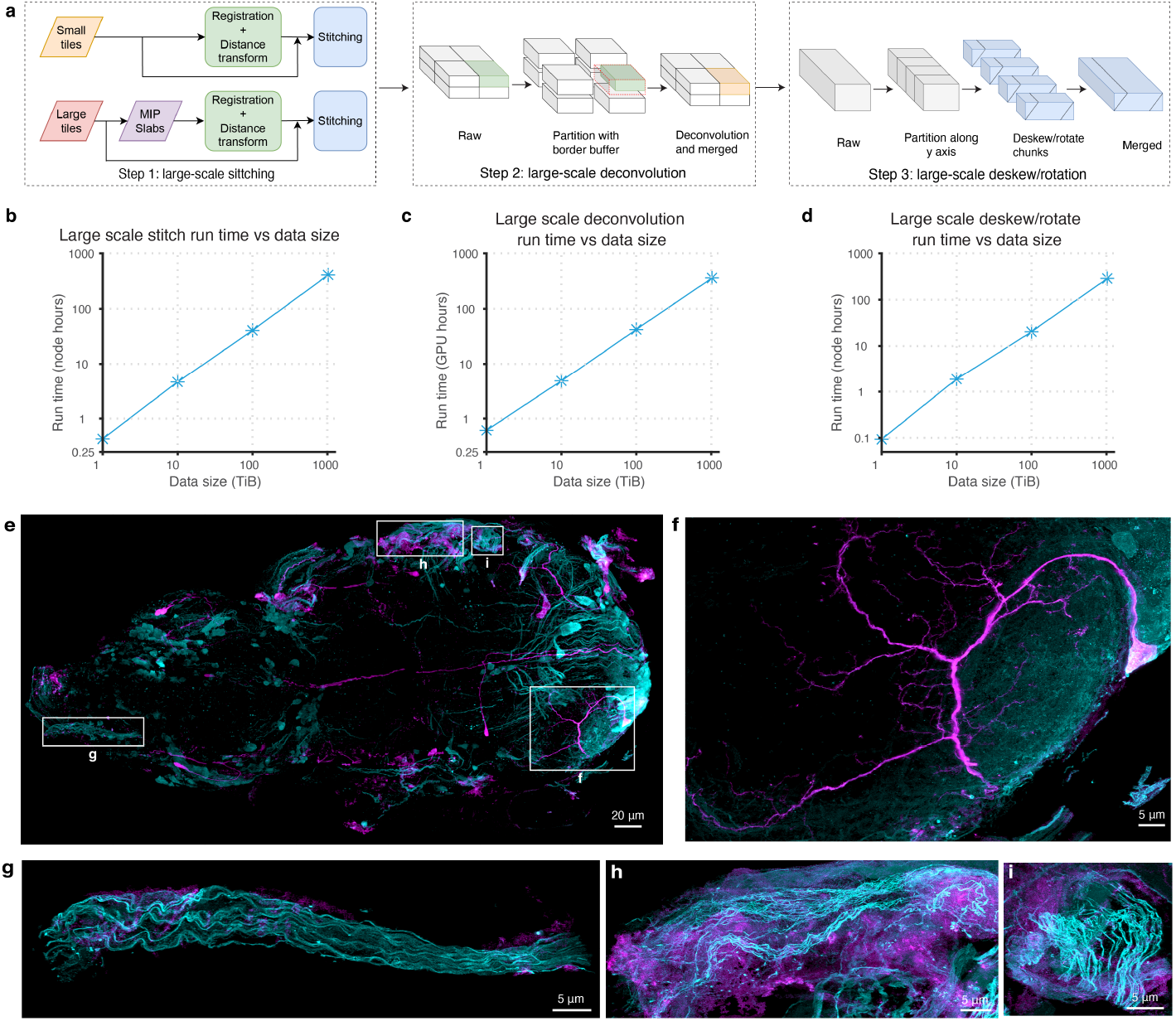
Large-scale processing. **a**, Schematic of processing steps for large-scale stitching, deconvolution, and deskew/rotation. For deconvolution, the data is split into subvolumes in all three axes with an overlap border size of slightly over half of the PSF size. For deskew and rotation, the data is split along the y-axis with a border of one slice. **b**, total run times for large-scale stitching of a single volume with size ranging from 1 TiB to 1 PiB. **c**, total run times for largescale deconvolution of a single volume with size ranging from 1 TiB to 1 PiB. **d**, total run times for large-scale deskew and rotation of a single volume with size ranging from 1 TiB to 1 PiB. Each benchmark in **b**-**d** was repeated three times independently. The resulting standard deviations are smaller than the data markers in the plots. **e**, MIP view of the entire fly VNC at 8× expansion. Cyan: VGlut^MI04979^-LexA::QFAD, and purple: MN-GAL4. Intensity *γ* = 0.5. **f** -**i**, MIP views of cropped regions from **e**. Intensity *γ* = 0.75 for all four regions.

When handling datasets that exceed memory capacity, certain processing steps become challenging. ZarrStitcher already enables stitching data that exceeds memory limitations as long as the intermediate steps can be fitted into memory. For stitching with even larger tiles, we developed a maximum intensity projection (MIP) slab-based stitching technique (Fig. 6a, left) where tiles are downsampled by different factors for different axes (e.g., 2× for the xy axes and 100× for z) to generate MIP slabs that fit into memory. These slabs are used to calculate registration information and estimate distance-based weights for feather blending, ensuring accurate stitching of the complete dataset (Fig. S7b, S7c-f bottom rows).

For deskew, rotation, and deconvolution, we distribute subvolumes of large data among multiple workers for faster processing and merge the results into the final output (Fig. 6a, middle and right). Zarr seamlessly enables this process.

In many imaging scenarios, a substantial amount of data beyond the boundary of the specimen is empty to ensure complete coverage. Processing these empty regions is unnecessarily inefficient, particularly for deconvolution (Fig. S10a-b, f). We therefore define the boundary based on MIPs across all three axes and skip the empty regions for large-scale deskew/rotation (Fig. S10c-e) and deconvolution (Fig. S10f-h).

With the above techniques, petabyte-scale processing becomes feasible and efficient. Processing time scales linearly for stitching, deconvolution, and deskew/rotation for data sizes ranging from 1 TiB to 1 PiB (Fig. 6b-d).

As an example, we processed a 38 TiB image volume of the Drosophila adult ventral nerve cord (VNC) at 8× expansion, as shown in Fig. 6e-i and Movie S5. All glutamatergic neurons, which include all motor neurons, are shown in cyan, and a subset of VNC neurons that includes a small number of these motor neurons is shown in purple. The ability to image, process, and visualize major complete anatomical regions such as the VNC at nanoscale resolution in multiple colors at such speeds opens the door to study the stereotypy and variability of neural circuits at high resolution over long distances, across large populations, different sexes, and multiple species.

## 3 Discussion

Our software achieves real-time processing at the multi-terabyte per hour acquisition rates of modern scientific cameras, for the extended times and/or large volumes that produce petabyte-scale data sets. It can be applied to many imaging modalities but includes deskew and rotation operations specifically useful in light sheet microscopy.

One limitation of the current pipeline is that it only supports rigid registration to compensate for sample translation, which performs well in most scenarios. However, it may not be suitable for multiview registration or fusion, image tiles with rotation, shrinking, swelling, or warping, which would require non-rigid methods such as elastic registration [44]. While zstd compression in Zarr is helpful, storing raw and intermediate data for petabyte-scale or larger datasets may still require hundreds of terabytes to petabytes of storage. Real-time preprocessing of raw data followed by massive compression during acquisition may be necessary to tackle this challenge.

Notably, our software is at least tenfold more efficient computationally than existing processing solutions, which can be used to either increase experimental throughput or decrease the number (and hence the cost) of computing nodes needed. In the former case, high throughput could prove useful in obtaining high-quality training data for deep learning image processing tasks [45–47], such as deconvolution [48], denoising [49], or registration [50]. The speed of the pipeline is also attractive for combining with multi-GPU 4D visualization [51] to monitor vast image-based biological experiments in real-time, including high throughput, high-resolution 3D drug screening [52], large tissue or whole organism spatial transcriptomics [53], or long-term imaging of subcellular dynamics in live multicellular organisms [54, 55].

## Online methods

### Generic computing framework

Our generic computing framework supports both single machines and large-scale Slurm-based computing clusters with CPU and/or GPU node configurations. The conductor job orchestrates the processing after it receives a collection of function strings (a MATLAB function call or a Bash command executed by each worker), input file names, output file names, and relevant parameters for job settings, such as required memory, the number of CPU cores, and the system environment. The conductor job initially checks for the presence of output files, skipping those that already exist. In single-machine setups or when Slurm job submission is disabled, the conductor job will sequentially execute tasks. Conversely, in cluster environments with the Slurm job scheduler, the conductor job formats and submits Slurm commands based on the function strings and job parameters, delegating tasks to workers in the cluster. It continuously monitors these jobs, ensuring the completion of the tasks. If a worker job fails, the conductor job resubmits it with an increased memory and CPU resources, often doubling the original specifications, until all tasks are completed, or a preset maximum retry limit is reached. Additionally, the framework allows users to define a custom configuration file. This feature tailors Slurm-related parameters to specific needs, ensuring adaptability to various user-defined function strings and compatibility with different Slurm-based computing clusters.

### Fast Tiff and Zarr readers and writers

Our Tiff reader/writer leverages the capabilities of the libtiff library in C++ with the MATLAB MEX interface. When reading, a binary search is used to determine the number of z slices by identifying the last valid slice, as there is no direct way to query the number of z slices in libtiff. The OpenMP framework is then used to distribute the reading tasks across multiple threads, partitioning the z slices into evenly sized batches (except for the last one). For large 2D images, the Tiff strips are partitioned to facilitate multi-threaded reading using the OpenMP framework. For the Tiff writer, LZW compression from libtiff is adapted to support compression on individual z slices. This approach enables parallel compression across z slices, leveraging the OpenMP framework for multi-threading. The final compressed data is written to disk using a single thread since a Tiff file is a single container, making parallel writing of compressed data infeasible.

As MATLAB lacks a native Zarr reader and writer, we developed custom C++ code that complies with the Zarr specification (version 2) with enhanced parallelization. This code is also integrated with MATLAB through the MEX interface. In our implementation, the OpenMP framework is used for both reading and writing to distribute the tasks across multiple threads, treating each chunk as a separate task. We use the compression algorithms from the Blosc library [56], which introduces an additional layer of multi-threading, thus optimizing the use of system resources. Zstd compression with a level 1 setting is used to achieve an optimal balance of compression ratio and read/write time. The high compression ratio of zstd substantially reduces the overall data size, reducing network load, particularly in extensive high-throughput processing scenarios where the network is often the primary bottleneck. By default, we read and write Zarr files in the “ Fortran” (column-major layout) order since MATLAB is based on “Fortran” order, and converting between “C” (row-major layout) and “Fortran” orders adds additional overhead.

### Combined deskew, rotation, resampling, and cropping

We execute deskew, rotate, and resampling (if needed) in a single step by combining these geometric transformations. The fundamental geometric transform involves:

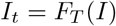

where *I* is the original image, *I*_*t*_ represents the transformed image, *F*_*T*_ () denotes the image warp function corresponding to the geometric transformation matrix *T*. The deskew operator applies a shear transformation defined by the shear transformation matrix *S*_*ds*_. In the rotation process, there are four sub-steps: translating the origin to the image center, resampling in the z-axis to achieve isotropic voxels, rotating along the y-axis, and translating the origin back to the starting index. Let the transformation matrices be denoted as *T*_1_, *S, R*, and *T*_2_, respectively. If resampling factors are provided (by default as 1), then there are three additional sub-steps in resampling: translating the origin to the image center, resampling based on the factors provided, and translating the origin back to the start index. Let the transformation matrices in these sub-steps be *T*_*R*1_, *S*_*R*_, and *T*_*R*2_, respectively.

Traditionally, these three steps are executed independently, resulting in multiple geometric transformations. However, this incurs substantial overhead in runtime and memory usage, particularly during the deskew step. Instead, we combine deskew, rotation, and resampling into one single step, resulting in a unified affine transformation matrix:

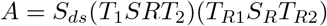

This affine transformation matrix can be directly applied to the raw image if the scan step size is sufficiently small. A quantity, denoted as “skew factor”, is defined to describe the relative step size as

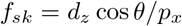

where *θ* ∈ (−*π/*2, *π/*2] is the skewed angle, *d*_*z*_ denotes the scan step size, and *p*_*x*_ is the pixel size in the xy plane. If *f*_*sk*_ ≤ 2, the direct combined processing operates smoothly without noticeable artifacts. For *f*_*sk*_ *>* 2, interpolation of the raw data within the skewed space is performed before deskew and rotation, taking account of the proper relative positions of slices. Neighboring slices above and below are utilized to interpolate a z slice. Let *w*_*s*_ and *w*_*t*_ = 1 − *w*_*s*_ represent the normalized distances (ranging from 0 to 1) along the z-axis between the neighboring slices and the target z slice. In the interpolation, we first create two planes aligned with the correct voxel positions of the target z slice by displacing the neighboring slices with a specific distance in the *x* direction (*w*_*s*_*d*_*z*_ cos *θ* and *w*_*t*_*d*_*z*_ cos *θ*, respectively). Following this, the target z slice is obtained by linearly interpolating these two planes along the z-axis with weights 1− *w*_*s*_ and 1 −*w*_*t*_. Because the image warp function permits the specification of the output view, we have also incorporated a cropping feature by providing a bounding box that allows us to skip empty regions or capture specific regions.

In acquisition modes where the deskew operation is unnecessary (e.g. objective scan mode of LLSM), the above processing can still be applied, provided *S*_*ds*_ is replaced with the identity matrix.

### Deconvolution

Richardson-Lucy (RL) deconvolution has the form of

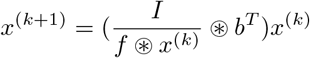

where *I* is the raw data, *f* is the forward projector (i.e., the point spread function (PSF)), *b* is the backward projector and *b*^*T*^ is the transpose of *b*, ⊛ denotes the convolution operator, and *x*^(*k*)^ is the deconvolution result in *k*-th iteration. In traditional RL deconvolution, *b* = *f*. In the OTF masked Wiener (OMW) method we use, the backward projector is generated with these steps:

1. The optical transfer function (OTF) *H* of the PSF *f* is computed, *H* = ℱ(*f*), where ℱ(·) represents the Fourier transform.
2. The OTF mask for the OTF support is segmented by applying a threshold to the amplitude |*H*|.The threshold value is determined by a specified percentile (90% by default) of the accumulated sum of sorted values in |*H*| from high to low.
3. The OTF mask undergoes a smoothing process, retaining only the central object, followed by convex hull filling. For deskewed space deconvolution, the three major components are kept after object smoothing and concatenated into a unified object along the z-axis, followed by convex hull filling.
4. The distance matrix *D* is computed with the image center as 0, and the edge of the support as 1 with the ray distance from the center to the border of the whole image.
5. The distance matrix *D* is used to calculate the weight matrix *W* with the Hann window function for apodization, as expressed by the following formula:

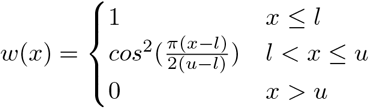

Where *l* and *u* are the lower and upper bounds for the relative distances. By default, *l* = 0.8 and *u* = 1 (edge of the support). For skewed space deconvolution, the weight matrix is given as a single distance matrix by adding the distance matrix from the corresponding three components together.
6. Calculate Wiener filter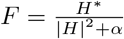, where *α* is the Wiener parameter, and *H*^∗^ denotes the conjugate transpose of *H*.
7. The backward projection in the Fourier space is expressed as *B* = *W* ⊙ *F*, where ⊙ denotes the Hadamard product operator (element-wise multiplication), and the backward projector in the real space is *b* = ℱ ^−1^(*B*), where ℱ ^−1^(·) represents the inverse Fourier transform.

The Fourier Shell Correlation (FSC) method [41] is used to determine the optimal number of traditional RL iterations and the optimal Wiener parameter in the OMW method. Here, the central portion of the volume, which is consistent in size across all three axes and covers sufficient content (i.e., 202×202×202 for a volume with size 230×210×202), is employed to compute the relative resolution. By default, the FSC is calculated with a radius of 10 pixels and an angle interval of *π/*12. Cutoff frequencies for relative resolution are determined using one-bit thresholding [57] by default, or can be user-defined. The relative resolution across iterations (or different Wiener parameters) is plotted. In [36], it was determined that a slightly higher threshold produced better results (Fig. S5g, purple circle). In practice, the optimal number of RL iterations or the Wiener parameter is defined by the value closest to 1.01 times the minimum of the curve beyond the point where the curve reaches its minimum value.

### Stitching

The stitching process requires a CSV meta-file documenting file names and corresponding coordinates. The pipeline consists of three steps: Tiff to Zarr conversion (or preprocessing), cross-correlation registration, and parallel block stitching (fusion). The overall stitching workflow is governed by a conductor job in the generic computing framework. For Tiff to Zarr conversion and/or processing on individual tiles, the conductor job distributes tasks to individual worker jobs, assigning one worker for each tile. Each worker: a) reads its data using the Cpp-Tiff or Cpp-Zarr (if existing Zarr data need rechunking or preprocessing) reader depending on the format; b) performs optional processing such as flipping, cropping, flat field correction, edge erosion, or other user-defined operations; and c) writes the processed data using the Cpp-Zarr writer.

Following file conversion, stitching can be executed directly using the input tile coordinates, or normalized cross-correlation registration [42] can be employed first to refine and optimize the coordinates before stitching. In the registration, the conductor job utilizes coordinate information and tile indices to establish tile grids and identify neighboring tiles with overlaps. Cross-correlation registration is performed for overlapping tiles that are direct neighbors, defined as those whose tile indices differ by 1, and only in one axis. To optimize computing time and memory usage, only the overlap regions for the tiles are loaded, including a buffer size determined by the maximum allowed shifts along the xyz axes within one tile. We can also downsample the overlap data to achieve faster cross-correlation computing. The optimal shift between the two tiles is identified as the one exhibiting the maximum correlation within the allowable shift limits. We include a feature to exclude shifts for pairs with the maximum correlation values below a user-defined threshold. After completing the cross-correlation computation for all pairs of direct neighbor tiles, we determine the shifts for all tiles using either a local or global method. The local approach is based on the concept of the minimum spanning tree, where the pairs of overlapping tiles are pruned to form a tree based on the correlation values from high to low, followed by registration with the pairwise optimal shifts. In the global approach, the optimal final shifts are calculated from the pairwise relative shifts through a nonlinear constrained optimization process:

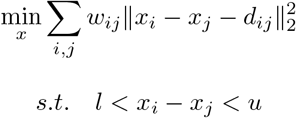

where **x**_*shift*_ ={*x*_1_, …, *x*_*n*_} are the final shifts for the tiles. *d* _*ij*_ is the pairwise relative shift between tile *i* and *j, w*_*ij*_ is the weight between tile *i* and *j* based on the squares of max cross-correlation values. *l* and *u* are the lower and upper bounds for the maximum allowable shift. The goal is to position all tiles at optimal coordinates by minimizing the weighted sum of the squared differences between their distances and the pairwise relative shifts while adhering to the specified maximum allowable shifts.

For images collected by subregions (batches) that have different tile grids, we employ the global method for tiles within each subregion. Subsequently, the subregions are treated as super nodes, and a nonlinear constrained optimization is applied to those nodes, by minimizing the sum of squared differences of the centroid distances to the averaged shift distances.

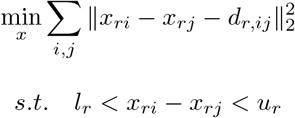

where *x*_*ri*_ and *x*_*rj*_ are the centroid coordinates for subregions *i* and *j, l*_*r*_ and *u*_*r*_ are lower and upper bounds for the maximum allowable shifts across subregions. The averaged shift distance, denoted as *d*_*r,ij*_, is determined by a weighted average of the absolute shifts across subregions, as expressed below

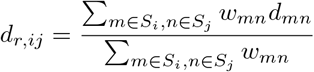

where *w*_*mn*_ is the cross-correlation value at the optimal shift between tiles *m* ∈ *S*_*i*_, and *n* ∈ *S*_*j*_, and *S*_*k*_ denotes the set of tiles in subregion *k*. Once the optimal shifts for the subregions are obtained, the last step is to reconstruct the optimal shifts for the tiles within each subregion by applying the optimal shifts of the centroid of the subregion to the coordinates of the tiles in it. The final optimal shifts are then applied to the tile coordinates to determine their final positions.

After registration, the conductor job determines the final stitched image size and the specific locations to place the tiles. To facilitate parallel stitching, the process is executed region by region in a non-overlapping manner. These regions are saved directly as one or more distinct chunk files in Zarr format. For each region, information about the tiles therein and their corresponding bounding boxes are stored. The conductor job submits stitching tasks to worker jobs. If the region comes from one tile, the data for the region is saved directly. If the region spans multiple tiles, these must be merged into a single cohesive region. For the overlap regions, several blending options are available: “none”, “mean”, “median”, “max” and “feather”. For the “none” option, half of the overlap region is taken from each tile. For the “mean”, “median”, and “max” options, the voxel values in the stitched region are calculated as the mean, median, and maximum values from the corresponding voxels in the overlapping regions, respectively. Feather blending involves calculating the weighted average across the tiles [43]. The weights are the power of the distance transform of the tiles as follows:

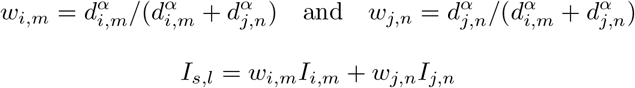

where *d*_*i,m*_ and *d*_*j,n*_ are distance transforms for voxel *m* in tile *i* and voxel *n* in tile *j, α* is the order (10 by default), *I*_*i*_ and *I*_*j*_ are the intensities from image *I*_*i*_ and *I*_*j*_, and *I*_*s*_ is the stitched image. Here we assume voxel *m* in tile *i* and voxel *n* in tile *j* are fused to voxel *l* in the stitched image. For the distance transform, we utilize a weighted approach, applying the distance transform to each z slice and then applying the Tukey window function across z slices to address the anisotropic properties of voxel sizes. When all tiles are the same size, we compute the weight matrix for a single tile and apply it across all other tiles in the stitching process to save computing time. The final stitched image is obtained once all the subvolumes are processed.

### Large-scale processing

For stitching involving large tiles where intermediate steps above exceed memory capacities, including large, stitched subregions, challenges arise in the registration and calculation of the distance transformation for feather blending, due to the need to load large regions or tiles into memory. In such cases, we use MIP slabs for the registration and distance transform. These are computed across all three axes with downsampling factors [*M*_*x*_, *M*_*y*_, *M*_*z*_, *m*_*x*_, *m*_*y*_, *m*_*z*_]. The MIP slab for each specific axis is computed using the major down sample factor *M*_*i*_ for that axis, and the minor down sampling factors *m*_*j*_ and *m*_*k*_ for the other two. To enrich the signal for cross-correlation in sparse specimens, we use max pooling, i.e., taking the max value in the neighborhood for the downsampling. Alternatively, we can also smooth the initial data by linear interpolation before max pooling. For the registration, normalized cross-correlation is calculated between direct neighbor tiles using all three MIP slabs, generating three sets of optimal shifts. The optimal shifts from the minor axes are then averaged to obtain the final optimal shifts, with weights assigned based on the squares of the cross-correlation values. For the distance transform, only the MIP slab along the z-axis (major axis) is employed to compute the weights for feather blending. In the stitching process, for overlapping regions, the down-sampled weight regions are up-sampled using linear interpolation to match the size of the regions in the stitching. The up-sampled weights are then utilized for feather blending, following the same approach as employed for stitching with smaller tiles.

For large-scale deskew and rotation, tasks are divided across the y-axis based on the size in the x and z axes, with a buffer of one or two pixels on both sides in the y-axis. These tasks are then allocated to individual worker jobs for processing, with the results saved as independent Zarr regions on disk. MIP masks can be used to define a tight boundary for the object to optimize efficiency in data reading, processing, and writing. We also perform deskewing and rotation for the MIP along the y-axis to define the bounding box for the output in the xz axes. The geometric transformation function directly relies on this bounding box to determine the output view to minimize the empty regions, thereby further optimizing processing time, memory, and storage requirements. For large-scale deconvolution, tasks are distributed across all three axes, ensuring that regions occupy entire chunk files. An additional buffer size, set to at least half of the PSF size (plus some extra size, 10 by default), is included to eliminate edge artifacts. MIP masks are again used to define a tight specimen boundary to speed computing. In a given task, all three MIP masks for the region are loaded and checked for empty ones. If a mask is empty, deconvolution is skipped, resulting in an output of zeros for that region.

### Image processing and simulations

All images were processed using PetaKit5D. Flat-field correction was applied for the large field of view cell data (Fig. 5f-h), phase contrast data (Fig. 5c-e), and VNC data (Fig. 6e-j) with either experimentally collected flat-fields or ones estimated based on the data using BaSiC software [58].

The images used to benchmark different readers and writers, deskew/rotation, and deconvolution algorithms were generated by cropping or replicating frames from a uint16 image of size 512 × 1, 800 × 3, 400. The stripped line patterns used to compare deconvolution methods were simulated using the methodology outlined in [36]. The confocal PSF for the given pinhole size used in the stripped line pattern simulation was generated based on the theoretical widefield PSF. We benchmarked large-scale stitching from 1 TiB to 1 PiB using one channel of the VNC dataset with 1,071 tiles, each sized at 320 × 1, 800 × 17, 001. The datasets were created by either including specific numbers of tiles or replicating tiles across all three axes based on the total data size from 1 TiB to 1 PiB, as specified in the table below. We benchmarked large-scale deconvolution and deskew/rotation using the stitched VNC dataset (15, 612 × 28, 478 × 21, 299, uint16) by either cropping or replicating the data in all three axes to generate the input datasets, as indicated in the table below.

|  | Stitching<br>(# tiles) | Deconvolution<br>(input size) | Deskew/rotation<br>(input size) |
| --- | --- | --- | --- |
| 1 TiB | 57 | $6,046 \times 11,027 \times 8,248$ | $2,722 \times 34,367 \times 5,879$ |
| 10 TiB | 562 | $13,024 \times 23,757 \times 17,769$ | $5,864 \times 74,041 \times 12,665$ |
| 100 TiB | 5,644 | $28,059 \times 51,183 \times 38,280$ | $12,632 \times 159,516 \times 27,284$ |
| 1 PiB | 57,392 | $60,931 \times 111,145 \times 83,127$ | $27,431 \times 346,393 \times 59,248$ |

The table above shows the number of tiles for stitching, and the input data dimensions for deconvolution and deskew/rotation in large-scale processing benchmarks.

### Computing infrastructures

Our computing cluster has 38 CPU/GPU computing nodes: 30 CPU nodes (24 nodes with dual Intel Xeon Gold 6146 CPUs, 6 nodes with dual Intel Xeon Gold 6342 CPUs) and 8 GPU nodes (3 nodes with dual Intel Xeon Silver 4210R and 4 NVIDIA Titan V GPUs each, 4 nodes with dual Intel Xeon Gold 6144 and 4 NVIDIA A100 GPUs each, and 1 NVIDIA DGX A100 with dual AMD EPYC 7742 CPUs and 8 NVIDIA A100 GPUs). The Intel Xeon Gold 6146 CPU and GPU nodes have 512 GB RAM on each node, the Intel Xeon Gold 6342 CPU nodes have 1024 GB RAM on each node, and the NVIDIA DGX A100 has 2 TB RAM. The hyperthreading on all Intel CPUs was disabled. Benchmarks were performed on hardware aged approximately three to four years. We have four flash data servers, including a 70 TB (SSD, Supermicro), two 300 TB (NVMe, Supermicro), and a 1000 TB parallel file system (VAST Data). We also accessed the Perlmutter supercomputer from the National Energy Research Scientific Computing Center (NERSC), with both CPU and GPU nodes. Each CPU node is equipped with two AMD EPYC 7713 CPUs and 512 GB RAM; each GPU node has a single AMD EPYC 7713 CPU, 4 NVIDIA A100 GPUs, and 256 or 512 GB RAM.

### Microscope hardware

Light sheet imaging was performed on a lattice light-sheet microscope comparable to a published system [54]. Two lasers, 488 nm and 560 nm (500 mW, MPB Communications 2RU-VFL-P-500-488-B1R, and 2RU-VFL-P-1000-560-B1R), were employed as the light sources. Water immersion excitation (EO, Thorlabs TL20X-MPL) and detection objectives (DO, Zeiss, 20×, 1.0 NA, 1.8 mm FWD, 421452-9800-000) were used for imaging with a sCMOS camera (Hamamatsu ORCA Fusion). The oblique illumination microscopy was also performed on the modified lattice light-sheet microscope using a 642 nm laser illuminated through the EO, and imaged using an inverted DO (Zeiss, 20×, 1.0 NA, 1.8 mm FWD, 421452-9880-000). Widefield and Confocal imaging were performed on an Andor BC43 Benchtop Confocal Microscope (Oxford Instruments) with a Nikon Plan Apo 40×, 1.25 NA SIL Silicone objective (Nikon, MRD73400), a 488 nm laser (Oxford Instruments, Andor Borealis), and a modified Andor Zyla sCMOS camera (Oxford Instruments, 4.1 MP, 6.5 *µ*m pixel size). Two-photon microscopy was performed on a custom-built microscope equipped with an upright DO (Zeiss, 20×, 1.0 NA, 1.8 mm FWD, 421452-9880-000), pulsed laser (Coherent, Chameleon LS), deformable mirror (ALPAO, DM69), and MPPC modules (Hamamatsu, C13366-3050GA and C14455-3050GA). The imaging conditions for the datasets can be found in Table S4.

### Cell culture and imaging

Pig kidney epithelial cells (LLC-PK1, a gift from M. Davidson at Florida State University) cells and HeLa cells were cultured in DMEM with GlutaMAX (Gibco, 10566016) supplemented with 10% fetal bovine serum (FBS; Seradigm) in an incubator with 5% CO_2_ at 37°C and 100% humidity. LLC-PK1 cells stably expressing the ER marker mEmerald-Calnexin and the chromosome marker mCherry-H2B were grown on coverslips (Thorlabs, CG15XH) coated with 200 nm diameter fluorescent beads (Invitrogen FluoSpheresTM Carboxylate-Modified Microspheres, 505/515 nm, F8811). When cells reached 30 − 80% confluency, they were imaged at 37°C in Leibovitz’s L-15 Medium without Phenol Red (Gibco catalog # 21-083-027), with 5% fetal bovine serum (ATCC SCRR-30-2020™), and an antibiotic cocktail consisting of 0.1% Ampicillin (ThermoFisher 611770250), 0.1% Kanamycin (ThermoFisher, 11815024) and 0.1% Penicillin/Streptomycin (ThermoFisher, 15070063). HeLa cells were cultivated on 25 mm coverslips until approximately 50% confluency was achieved. They were imaged in the same media as above.

### Mouse brain sample preparation and imaging

All mice experiments were conducted at Janelia Research Campus, Howard Hughes Medical Institute (HHMI) in accordance with the US National Institutes of Health Guide for the Care and Use of Laboratory Animals. Procedures and protocols were approved by the Institutional Animal Care and Use Committee of the Janelia Research Campus, HHMI.

Transgenic Thy1-YFP-H mice of 8 weeks or older with cytosolic expression of yellow fluorescent protein (YFP) at high levels in motor, sensory, and subsets of central nervous system neurons were anesthetized with isoflurane (1− 2% by volume in oxygen) and placed on a heated blanket. An incision was made on the scalp followed by removing of the exposed skull. A cranial window made of a single 170-um-thick coverslip was embedded in the craniotomy. The cranial window and a headbar were sealed in place with dental cement for subsequent imaging. A direct wavefront sensing method [59] was used for adaptive optical correction prior to image acquisition. Aberrations at each volumetric tile were independently measured and corrected using a pupil conjugated deformable mirror, and imaged at 16 Hz using Hamamatsu MPPC modules.

### Fly VNC sample preparation and imaging

A genetically modified strain of fruit flies (Drosophila melanogaster) was raised on a standard cornmeal-agar-based medium in a controlled environment of 25°C on a 12:12 light/dark cycle. On the day of eclosion, female flies were collected, and group housed for 4-6 days. The genotype was VGlut^MI04979^-LexA:QFAD/MN-GAL4 (attp40); 13XLexAop-Syn21-mScarle [JK65C], 20XUAS-Syn21-GFP [attp2]/MN-GAL4 [attp2] [60, 61]. Dissection and immunohistochemistry of the fly VNC were performed following the protocol in [62] with minor modifications. The primary antibodies were chicken anti-GFP (1:1000, Abcam, ab13970) and rabbit anti-dsRed (1:1000, Takara Bio, 632496). The secondary antibodies were goat anti-chicken IgY Alexa Fluor 488 (1:500, Invitrogen, A11039) and goat anti-rabbit IgG Alexa Fluor 568 (1:500, Invitrogen, A11011). VNC samples were prepared for 8× expansion as described in [62]. The imaging protocol for the expanded VNC sample was identical to that described in [12].

### Visualization and software

Movies were made with Imaris (Oxford Instruments), Fiji [33], Amira (Fisher Scientific), NVIDIA IndeX (NVIDIA), and MATLAB R2023a (MathWorks) software. Figures were made with MATLAB R2023a (MathWorks). Python (3.8.8) with Zarr-Python (2.16.1), tifffile (2023.7.10), TensorStore (0.1.45), pyclesperanto-prototype (0.24.2), and clij2-fft (0.26) libraries were used for benchmarking image readers and writers, deskew and rotation, and deconvolution. The traditional RL deconvolution method is an accelerated version of the original RL algorithm [39, 63]. It was implemented and adapted from MATLAB’s *deconvlucy*.*m* with enhancements such as GPU computing and customized parameters. Backward projectors for the WB deconvolution method were generated using the code from https://github.com/eguomin/regDeconProject. Spark versions of BigStitcher (https://github.com/JaneliaSciComp/BigStitcher-Spark) and https://github.com/saalfeldlab/stitching-spark were used for the stitching comparison. NVIDIA IndeX can be obtained from https://developer.nvidia.com/index with a free license for non-commercial research and education.

## Acknowledgments

We thank Dr. Lin He for providing access to the confocal microscope. We thank Jonathan Lefman and the NVIDIA IndeX team for sharing the NVIDIA Index software. We thank John White for managing our computing cluster. We gratefully acknowledge the support of this work by the Laboratory Directed Research and Development (LDRD) Program of Lawrence Berkeley National Laboratory under US Department of Energy contract No. DE-AC02-05CH11231. This research used resources of the National Energy Research Scientific Computing Center (NERSC), a U.S. Department of Energy Office of Science User Facility located at Lawrence Berkeley National Laboratory, operated under Contract No. DE-AC02-05CH11231 using NERSC award DDR-ERCAP0025501.

## Authors contributions

E.B. and S.U. supervised the project. X.R. wrote the manuscript with input from all coauthors. E.B. and S.U. edited the initial draft. X.R. designed the algorithms and implemented the software. M.M. implemented the fast image readers and writers, the Parallel Fiji Visualizer plugin, the Imaris file converter, and the Python wrappers under X.R.’s guidance. M.M. designed and developed the graphical user interfaces, and C.Y.A.H. contributed to the implementation. J.L.L. prepared the VNC sample and G.L. performed the imaging experiments. W.H. and A.N.K. prepared the cultured LLC-PK1 cells and F.G. performed the live cell imaging experiment. T.F. performed the imaging experiments for the live mouse brain imaging and phase imaging of HeLa cells. D.M. developed the microscope software for the imaging experiments. A.K. and M.N. helped set up the workflows for the real-time visualization movie using NVIDIA IndeX. J.L.H. prepared the sample and J.G.C performed the widefield and confocal imaging experiments for the LLC-PK1 cells. X.R. performed all image processing and analysis and made the figures. X.R. and M.M. made all movies with S.U.’s input.

## Data availability

The full datasets for this manuscript exceed the size limits of any data repository, but they will be shared upon reasonable request. The representative subsets of the full datasets can be downloaded from https://doi.org/10.5061/dryad.kh18932g4 and https://doi.org/10.5061/dryad.jq2bvq8jd.

## Code availability

The source code of the software is available at https://github.com/abcucberkeley/PetaKit5D. The version associated with this manuscript is https://doi.org/10.5281/zenodo.12661251. We also provide a Python version for the wrapper of the deployed version of PetaKit5D at https://github.com/abcucberkeley/PyPetaKit5D. The graphical user interface for the software can be downloaded from https://github.com/abcucberkeley/PetaKit5D-GUI. The Parallel Fiji Visualizer plugin can be accessed from GitHub (https://github.com/abcucberkeley/Parallel_Fiji_Visualizer) or Zenodo (https://doi.org/10.5281/zenodo.7613228). The code for replicating the benchmark results is available for download on Zenodo (https://doi.org/10.5281/zenodo.12676473). The Nvidia IndeX software can be acquired by following the instructions in Supplementary Note 5. Example code and data for data format conversion and visualization are available for download from Zenodo (https://doi.org/10.5281/zenodo.12539579).

