## Supplemental Information for "Image processing tools for petabyte-scale light sheet microscopy data"

[1]Department of Molecular and Cell Biology, University of California, Berkeley, CA 94720.
[2]Howard Hughes Medical Institute, Berkeley, CA 94720.
[3]Janelia Research Campus, Howard Hughes Medical Institute, Ashburn, VA 20417.
[4]NVIDIA, Berlin, 10623, Germany.
[5]Department of Physics, Helen Wills Neuroscience Institute, University of California, Berkeley, CA 94720.
[6]Molecular Biophysics and Integrated Bioimaging Division, Lawrence Berkeley National Laboratory, Berkeley, CA 94720.
[7]Chan Zuckerberg Biohub, San Francisco, CA 94158.
[8]Current address: Department of Microsystems Engineering, University of Freiburg, Freiburg, 79110, Germany.
[9]Current address: Department of Electrical and Computer Engineering, Princeton University, Princeton, NJ 08540.

The PDF file includes:

- Supplementary notes 1-5
- Supplementary figures S1-S12
- Supplementary tables S1-S4
- Legends of supplementary movies S1-S6

### Supplementary Notes

#### 1. Useful tools

PetaKit5D software incorporates several commonly used functions including cropping, resampling, max intensity projection (MIP) generation, Fourier Shell Correlation (FSC) analysis, Fast Fourier transform (FFT) analysis, point spread function (PSF) and optical transfer function (OTF) visualization, Tiff/Zarr conversion, visualization of segmented OTF, Imaris (Oxford Instruments) file converter, and Image List generator for ZarrStitcher. For large-scale cropping, resampling, and MIP generation, we adopt a chunk-based approach akin to that used for deconvolution and deskew/rotation. Here, a large Zarr image is divided into smaller overlapping subvolumes, each handled by individual workers. The overlap for subvolumes is unused in functions such as cropping and MIP generator. FSC analysis helps determine the number of iterations necessary for traditional RL methods or the Wiener parameter in the OMW method. FFT analysis calculates the 3D Fourier spectra and displays the principal planes along xy, xz, and yz for a given volume. PSF and OTF visualization tools aid in comparing the empirical data and theoretical models to characterize the performance of the microscopes. The Tiff/Zarr converters allow for data format conversions. The OTF segmentation visualization tool helps determine the optimal parameters to segment the OTF for the OMW deconvolution method. The Imaris file converter combines our fast Tiff or Zarr readers with the ImarisWriter [1] to efficiently convert files into Imaris 10.0 compatible format. Lastly, the Image List generator creates a metadata CSV file for each dataset for ZarrStitcher using tile coordinate files (CSV), SQLite databases with tile coordinate information, or user-provided tile overlaps.

#### 2. Graphical user interface

We developed an easy-to-use interface for the PetaKit5D software, designed for users with little programming experience. The GUI was implemented with the Qt framework (The Qt Company) using C++ for cross-platform compatibility. With the GUI, users can easily execute processing steps interactively, including deskew/rotation, deconvolution, and stitching, simply by clicking buttons or filling in forms to input parameters for each step (Fig S11a). The GUI includes a main menu for browsing data paths, choosing processing steps, and setting common imaging and processing parameters. The GUI also allows users to use a custom configuration file for submitting Slurm jobs. Users can then navigate to the menus for the selected processing steps to enter processing-specific parameters. The final page allows users to start sequential processing locally or submit all jobs to a computing cluster.

The GUI also includes the set of useful tools such as cropping, Tiff/Zarr conversion, MIP generation, resampling, visualization of PSF and OTF, OTF segmentation, Imaris conversion, and Image List generation for ZarrStitcher (Fig. S11b, Supplementary Note 1).

We used MATLAB Compiler (MCC) to develop a deployed version of PetaKit5D for users without a MATLAB license. This version utilizes MATLAB runtime to execute essential functions. The GUI supports both MATLAB and deployed versions.

#### 3. Parallel Fiji Visualizer plugin

To streamline image opening within Fiji, we developed a dedicated Fiji plugin leveraging our fast Tiff/Zarr readers. This plugin presents a user-friendly drag-and-drop window (Fig S12a) to easily open Tiff or Zarr images, which utilizes the Cpp-Tiff or Cpp-Zarr libraries for efficient reading and subsequently display in Fiji. Our plugin significantly outperforms the native Fiji reader, with a speed improvement of 12-17× and 1.3-3.2× for Tiff and Zarr, respectively (Fig. S12b and c, Movie S6). The Cpp-Zarr reader benchmarks account for the additional overhead to flip the x and y axes in the Zarr opener (not currently supported by the N5/Zarr reader [2]) to make the axes consistent with the Tiff reader.

#### 4. Variations of Richardson Lucy deconvolution

Several variants of Richardson Lucy (RL) deconvolution techniques have been developed to address specific limitations of the original algorithm (the native method), including Biggs (referenced as the traditional approach in the manuscript) [3], total variation (TV) [4], non-circulant [5], Wiener-Butterworth (WB) [6], OTF-masked Wiener (OMW, this work) among others.

The Biggs method accelerates the iteration process by calculating an acceleration factor at each iteration using line search methods, resulting in speeds up to ten times faster than the native RL method (Fig. S6). Total variation (TV) RL deconvolution suppresses noise amplification by incorporating total variation regularization in the iteration process. The performance of the TV-RL method depends on the regularization factor and typically requires a similar number of iterations as the native RL method.

The non-circulant RL method is primarily used to reduce edge artifacts. It achieves this by padding the image with an empty region the size of the PSF and applying a weighted window function. This eliminates circulant artifacts at the image edges caused by FFT convolution and requires a similar number of iterations as the native RL method (if implemented based on the native method). However, it may significantly increase the processing time and the system or GPU memory requirements due to the need for padding data with the size of the PSF, especially for 3D images. If objects are fully contained within and away from the bounding box of the image, or if sufficient overlap is included for batch processing (e.g., large-scale processing for deconvolution described in the manuscript), the benefits of non-circulant deconvolution are negligible (Fig. S6).

WB and OMW are mainly designed to speed up the iteration processes by optimizing the backward projectors. Both approaches require significantly fewer iterations to converge compared to the Biggs, native, and non-circulant methods (Fig. S6). Since Biggs, TV, non-circulant, and our methods enhance RL deconvolution in different ways, the principles of these methods may be combined to further improve overall deconvolution performance.

#### 5. Real-time visualization using NVIDIA IndeX

We can couple the real-time processing capabilities (Table S3) with the NVIDIA IndeX software platform [7, 8] to leverage computing and GPU resources across multiple nodes for large-scale 5D data visualization, spanning hundreds of terabytes at full resolution. This platform offers different methods for real-time visualization using processed outputs from PetaKit5D.

The first approach converts data from Zarr to NanoVDB format. NanoVDB format is optimized for GPU and memory efficiency [9] and is particularly suited for visualizing sparse and compressed datasets that are common in microscopy. This approach uses GPUdirect storage that distributes data across multiple drives, each mapped to a GPU in systems like NVIDIA DGX A100, enabling fast data loading and rendering. Currently, the conversion process to NanoVDB can be a rate-limiting step since existing implementations are not optimized and the rendering requires specialized hardware/software configurations with GPUdirect support.

A second approach involves converting data to uncompressed raw format, then loading it into RAM for GPU access and caching for real-time rendering. This approach is compatible with standard multi-node, multi-GPU setups and does not require specialized hardware/software configurations. The limitation is the RAM capacity, typically in the terabyte range, restricting the amount of data that can be cached for real-time interactive rendering. The RAM and GPU requirements for larger datasets scale almost linearly with size to maintain the ability for real-time interactive visualization. Future improvements may include using a fast Zarr reader to directly access the Zarr files for real-time multi-GPU rendering with standard multi-node, or GPUdirect storage configurations.

##### 5.1 Deployment, execution, and availability of NVIDIA IndeX

NVIDIA IndeX (https://developer.nvidia.com/index) divides the three-dimensional data space into smaller portions, called subregions, for data-parallel visualization. The assigned GPUs in a system, such as those reserved in a supercomputer, process the subregions in parallel. In the present use case, visualization for the processed outputs from PetaKit5D requires a dedicated preparation and distribution scheme, as discussed below.

A set of Zarr files stores the processed microscopy data. Each file represents one channel and a time point, with the data divided into smaller chunks within each file. Let $S$ be the set of Zarr files and $D$ the set of subregions representing the data distribution for later visualization. A pre-processing step produces the subset for each subregion as follows:

```
for each d ∈ D:
   Determine a set S′ ⊂ S that includes all Zarr files that overlap with d
   for each s′ ∈ S′:
      Load portion that overlaps with d
      Store portion in d as a dense volume // Dense volume without any
          compression and sparsity
   end for

   // Data size reduction by exploiting homogeneous and empty space
   Encode sparsity:
      1) Determine homogeneous regions inside the volume where all values are
          equal
      2) Determine void, i.e., regions inside the volume where all values are
          equal to a user-defined threshold value that represents emptiness //
          Optional and user-defined

   // Data size reduction using optional quantization
   Apply data quantization, e.g., to 8-bit // Optional and user-defined

   // Data export in NanoVDB-formatted form
   Store volume data for subregion d
end for
```

To test the deployment and execution of NVIDIA IndeX, we included an example dataset and demo scripts for data conversion to either NanoVDB or raw binary formats (https://doi.org/10.5281/zenodo.12539579). When NanoVDB (https://developer.nvidia.com/nanovdb or https://developer.nvidia.com/blog/accelerating-openvdb-on-gpus-with-nanovdb) format is used for storing the volume subset data per subregion, it allows for encoding sparsity and quantization information (e.g. convert_zarr_to_raw.py and convert_raw_to_nanovdb.sh from Zenodo https://doi.org/10.5281/zenodo.12539579).

The resulting data distribution $D$ serves as input for the scalable visualization. NVIDIA IndeX supports several modes of operation for data distribution. It can either automatically create a data distribution scheme at the start while, for instance, leveraging the sparsity inside datasets, or it can adopt a predefined data distribution like the one given by $D$.

Launching NVIDIA IndeX with the data distribution on a computing cluster using the Slurm job scheduler[1] triggers the following steps automatically:

(1) Starts an NVIDIA IndeX instance on each allocated machine,
(2) Assign each subregion $d \in D$ evenly across the GPUs of the allocated machines,
(3) Importing the NanoVDB data for each subregion $d$ in parallel,
(4) Synthesizing images $f$ by rendering the subregions in parallel,
(5) When playing a different time point, repeat steps 3 and 4.

Once started, users can open a web browser and visit a URL connected to an NVIDIA IndeX web server (initialized in step 1 above). The web server provides a user interface to adjust the visual appearance of the dataset renderings and a live video stream for real-time interactions with the scene. Users can zoom, rotate, and explore different time points to inspect the data on the fly (e.g., Movie S3).

[1]NVIDIA IndeX is agnostic to the job scheduling and target systems.

Instead of using the NanoVDB data format, NVIDIA IndeX can visualize data directly from the raw binary format. This option is only suitable for smaller volumes that can fit into GPU memory without compression. The conversion from the Zarr format to the raw binary format can be done using a Python script (e.g., convert_zarr_to_raw.py from Zenodo https://doi.org/10.5281/zenodo.12539579) and is based on the basic NumPy array exporting from Python. Once the dataset is exported, the data can be loaded and visualized. If sufficient GPU memory is available, multiple raw binary timesteps can be preloaded and cached into GPU memory to facilitate rapid replay. This option has extended data loading time onto GPU memory and can leverage the combined VRAM when multiple GPUs are available.

NVIDIA IndeX comes with a free license for non-commercial research and education. For commercial use, please contact. The NVIDIA IndeX package can be requested through the following form: https://developer.nvidia.com/index-contact. The software package includes libraries for Linux, Windows, and Arm systems. It can be launched from within the package, scheduled as a Docker container, or integrated into a software distribution such as Paraview through a general C++ API (https://raytracing-docs.nvidia.com/nvindex/index.html). NVIDIA IndeX has been deployed in various settings, including supercomputers like Perlmutter at the National Energy Research Scientific Computing Center, on-premises data centers, the cloud, workstations, and laptops. The NVIDIA IndeX packages include getting-started information, a tutorial, and an extensible viewer for users to visualize their own data easily.

### Supplementary Figures

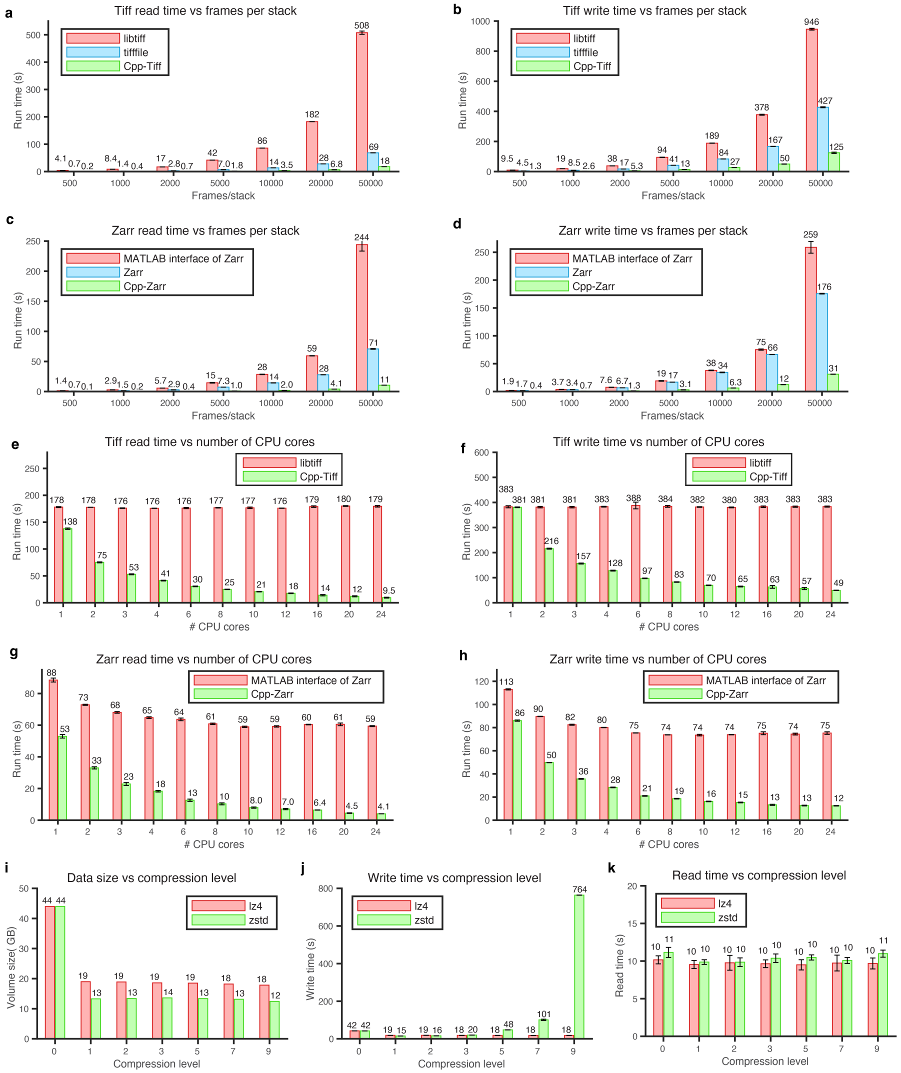


**Fig. S1**: Benchmarks for Tiff and Zarr readers and writers. **a-b**, run times of Tiff readers and writers for libtiff (MATLAB), tifffile (Python), and Cpp-tiff versus the number of frames. **c-d**, run times of Zarr readers and writers, comparing the MATLAB interface of Zarr, native Zarr (Python), and Cpp-Zarr across different numbers of frames. **e-f**, run times of Tiff readers and writers for libtiff (MATLAB) and Cpp-Tiff versus the number of CPU cores for an image stack of size $512 \times 1,800 \times 20,000$. **g-h**, run times of Zarr readers and writers for the MATLAB interface of Zarr, and Cpp-Zarr versus the number of CPU cores for an image stack of size $512 \times 1,800 \times 20,000$. **i-k**, data size and read/write times versus compression level for lz4 and zstd compressors. The benchmarks were run ten times independently on a 24-core CPU computing node (dual Intel Xeon Gold 6146 CPUs). All 24 cores were allocated for **a-d** and **i-k**, and varying numbers of CPU cores were allocated for **e-h**.

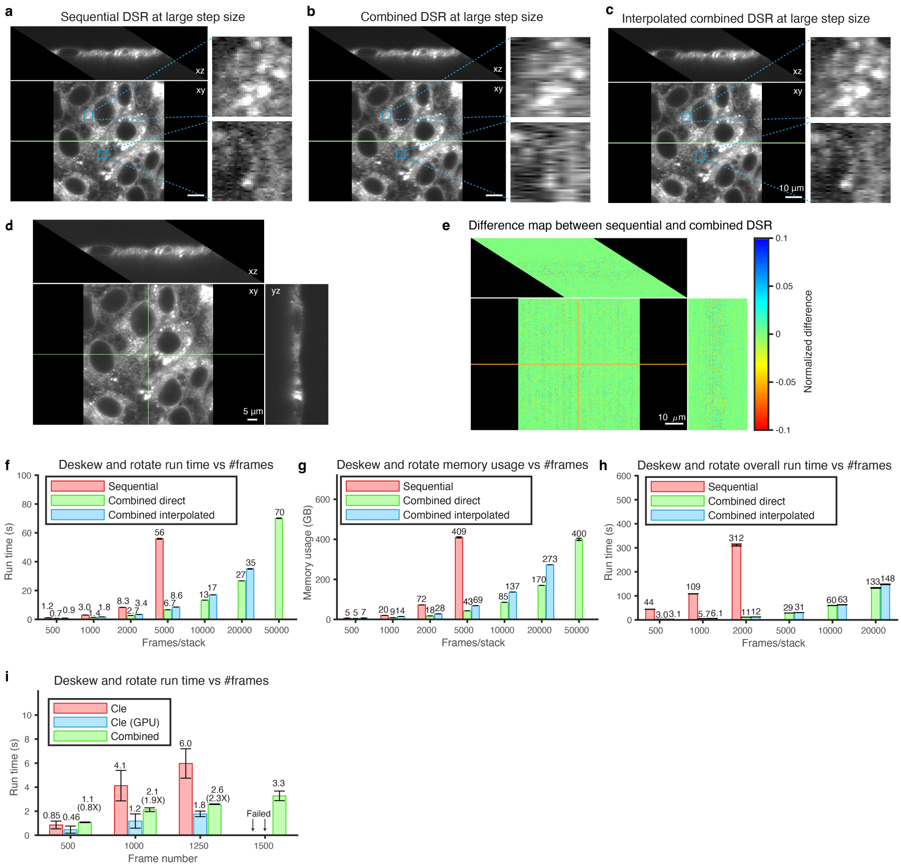


**Fig. S2**: Deskew and rotation. **a** sequential deskew/rotation for an image stack with a step size 0.6 $\mu$m between planes. **b-c**, combined deskew/rotation and interpolation plus combined deskew/rotation for the same image as in **a**. **d**, orthogonal views of combined deskew/rotation for the image in Fig. 3b. **e**, difference map between sequential and combined deskew/rotation for the images in Figs. 3b and S2d. **f-h**, benchmarks of sequential, combined, and interpolated combined deskew/rotation versus the number of frames, for run time (**f**), memory usage (**g**), and overall run time including reading and writing (**h**). **i**, benchmarks of the combined deskew/rotation method implemented in pyclesperanto using both CPU ("Cle") and GPU ("Cle (GPU)") and our combined interpolated approach ("Combined"). The method in pyclesperanto failed for images with 1,500 or more frames. For **f-i**, all images have a 32-bit frame size of $512 \times 1,800$ (xy). Each benchmark was run independently ten times on a 24-core CPU computing node (dual Intel Xeon Gold 6146 CPUs), except for "Cle (GPU)" which was run on a GPU node with 80 GB A100 GPUs.

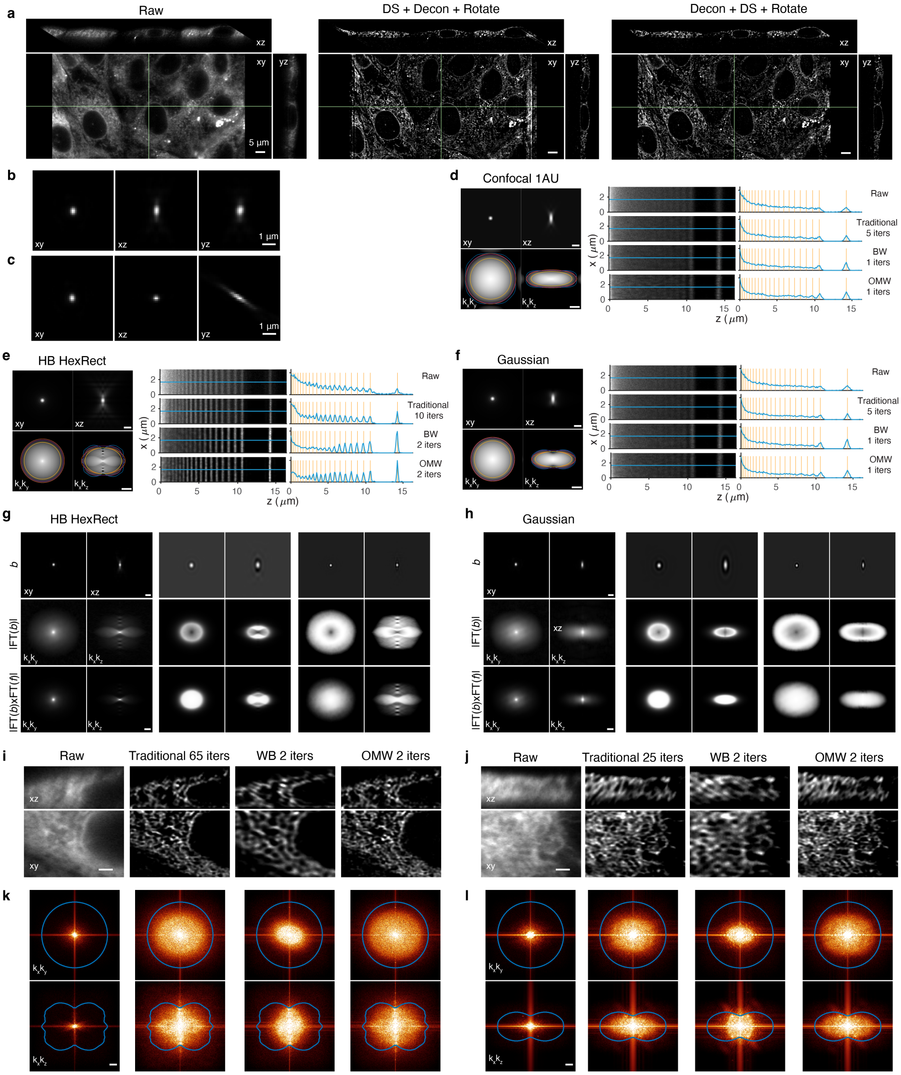


**Fig. S3**: Comparison of deconvolution methods for different light sheets. **a**, comparison of first deskew then deconvolution and rotation (center) versus first deconvolution then deskew and rotation. **b-c**, microscope PSF as seen in deskewed space (top) and skewed space (bottom). **d-f**, comparison of deconvolution of a simulated stripe pattern with different deconvolution methods for different light sheets. **g-h**, illustration of backward projectors for different deconvolution methods for harmonic-balanced (HB) HexRect, and Gaussian light sheets. **i-j**, comparison of cell images deconvolved by different deconvolution methods for HB HexRect and Gaussian light sheets. **k, l**, Fourier spectra for the raw and deconvolved images in **i** and **j**.

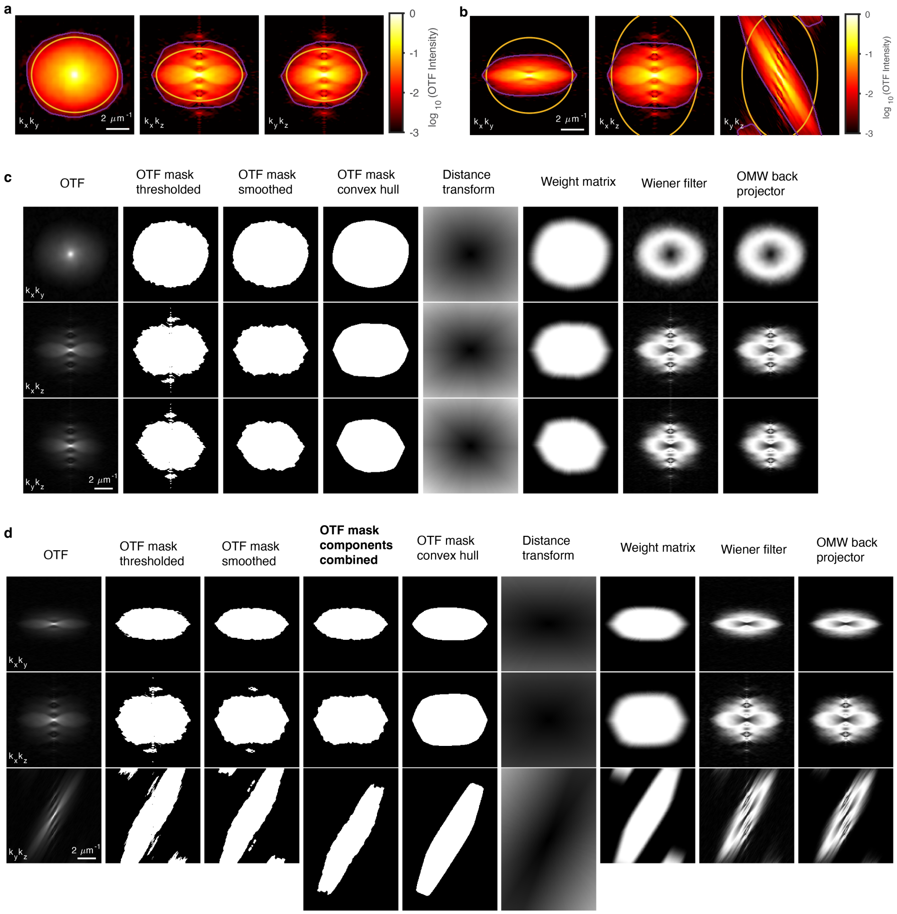


**Fig. S4**: Generation of the OMW backward projector. **a** and **b**, OTF support for WB (orange) and OMW (purple) methods for PSFs in deskewed or skewed space. **c**, OMW backward projector generation process using the PSF in the deskewed space. **c**, OMW backward projector generation process using the PSF in the skewed space.

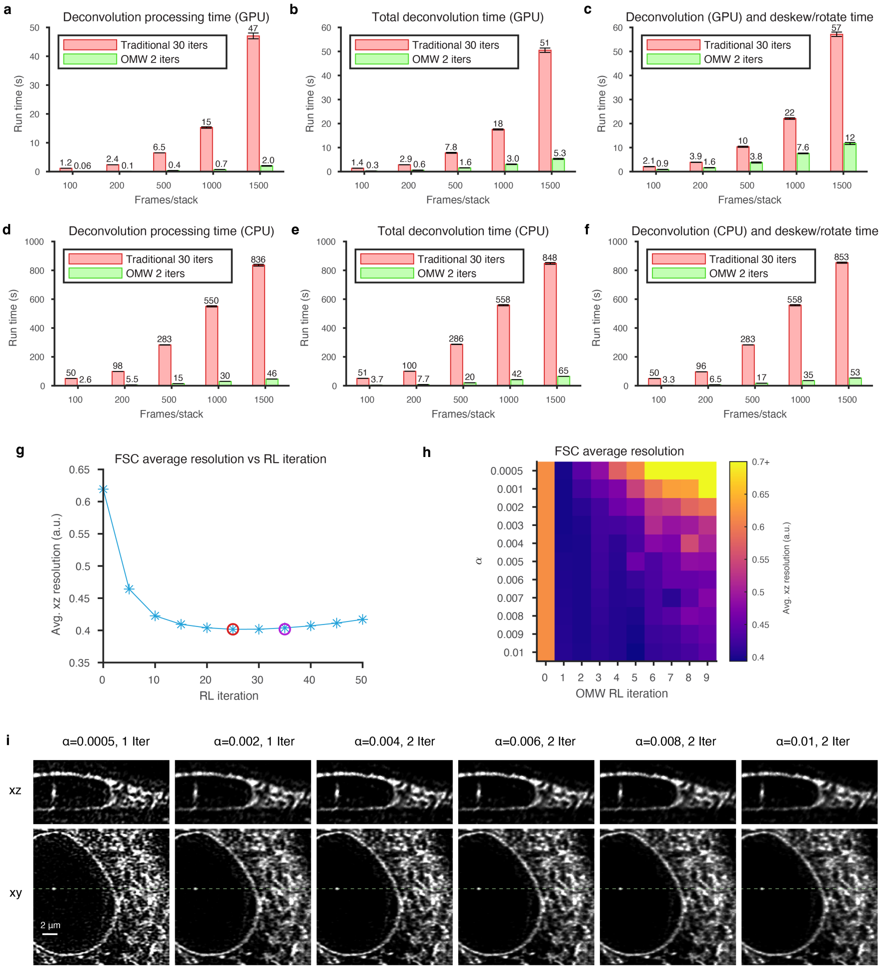


**Fig. S5**: Benchmarks comparing conventional to OMW deconvolution. **a, d**, deconvolution processing time along on GPU and CPU. **b, e**, deconvolution plus read/write time on GPU and CPU. **c, f**, deconvolution plus combined deskew/rotation time on GPU and CPU. **g**, Using FSC to pick the optimal number of iterations for deconvolution by the traditional RL method. **h**, Using FSC to select the Wiener $\alpha$ and the number of iterations for OMW deconvolution. **i**, Comparative images of OMW deconvolution with 1 or 2 iterations and four different values Wiener $\alpha$.

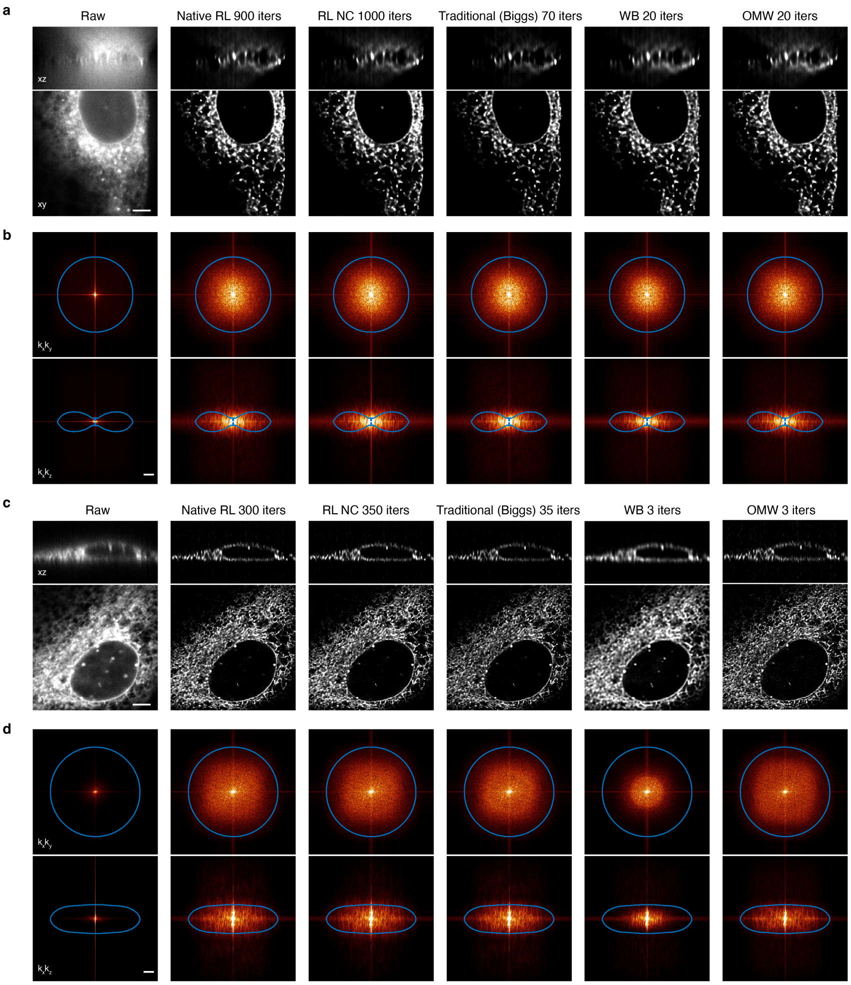


**Fig. S6**: Deconvolution methods comparison for widefield and confocal microscopy images. **a**, comparison of deconvolved orthoslices (scale bar: 5 $\mu m$) and **b**, Fourier spectra outputs for a widefield image (intensity $\gamma = 0.5$, scale bar: $1\,\mu m^{-1}$). **c**, comparison of deconvolved orthoslices (scale bar: 5 $\mu m$) and **d**, Fourier spectra outputs for a confocal image (intensity $\gamma = 0.5$, scale bar: $1\,\mu m^{-1}$). "RL NC" stands for the non-circulant RL method. The number of iterations is noted in the title for each method. For **b** and **d**, the blue lines indicate the theoretical OTF support. For **d**, the OTF support for 1 Airy unit (AU) is shown.

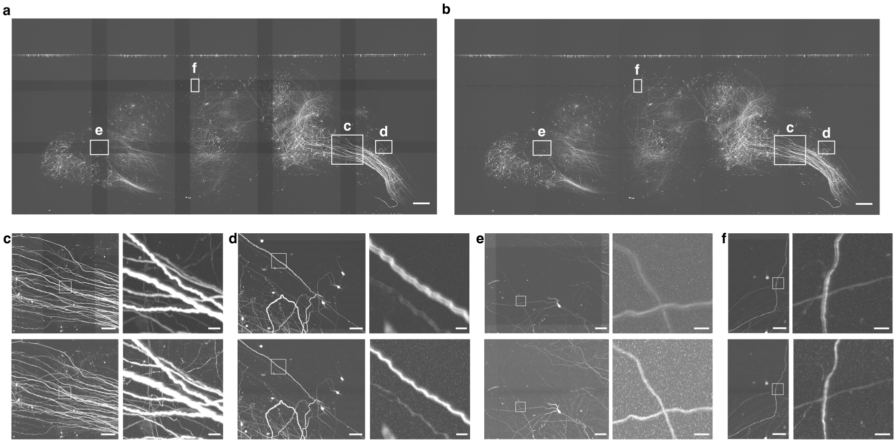


**Fig. S7**: Comparison of stitching quality between BigStitcher and ZarrStitcher for the whole mouse brain ExA-SPIM images with 4× expansion (Glaser et al. 2023). **a-b**, complete stitched brains with BigStitcher (**a**) and ZarrStitcher (**b**). **c-f**, expanded views of the boxed regions in **a-b** for BigStitcher (top rows) and ZarrStitcher (bottom rows). BigStitcher used the registration information from ZarrStitcher.

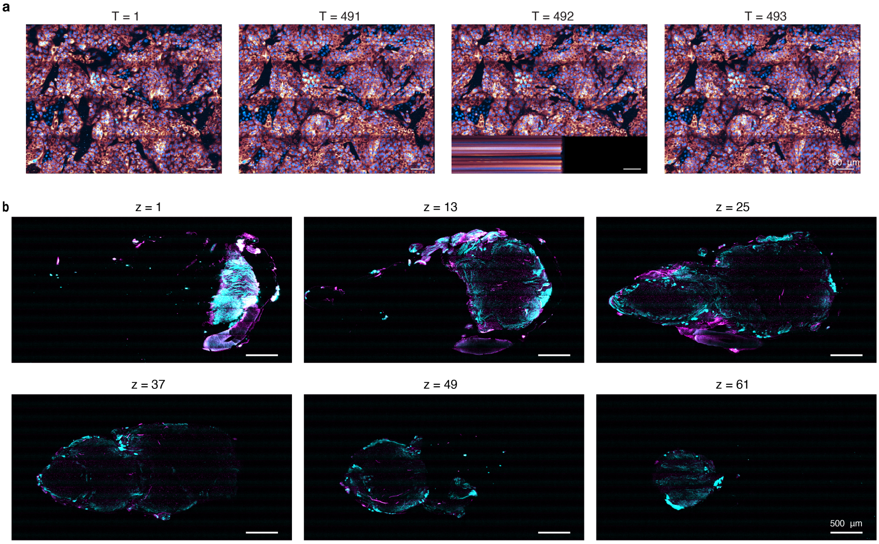


**Fig. S8**: Advantages of real-time processing and feedback for image acquisition. **a**, real-time identification of an acquisition error during large field-of-view live cell imaging. **b**, real-time images of stitched specimens across z layers to identify when and where to finish image acquisition.

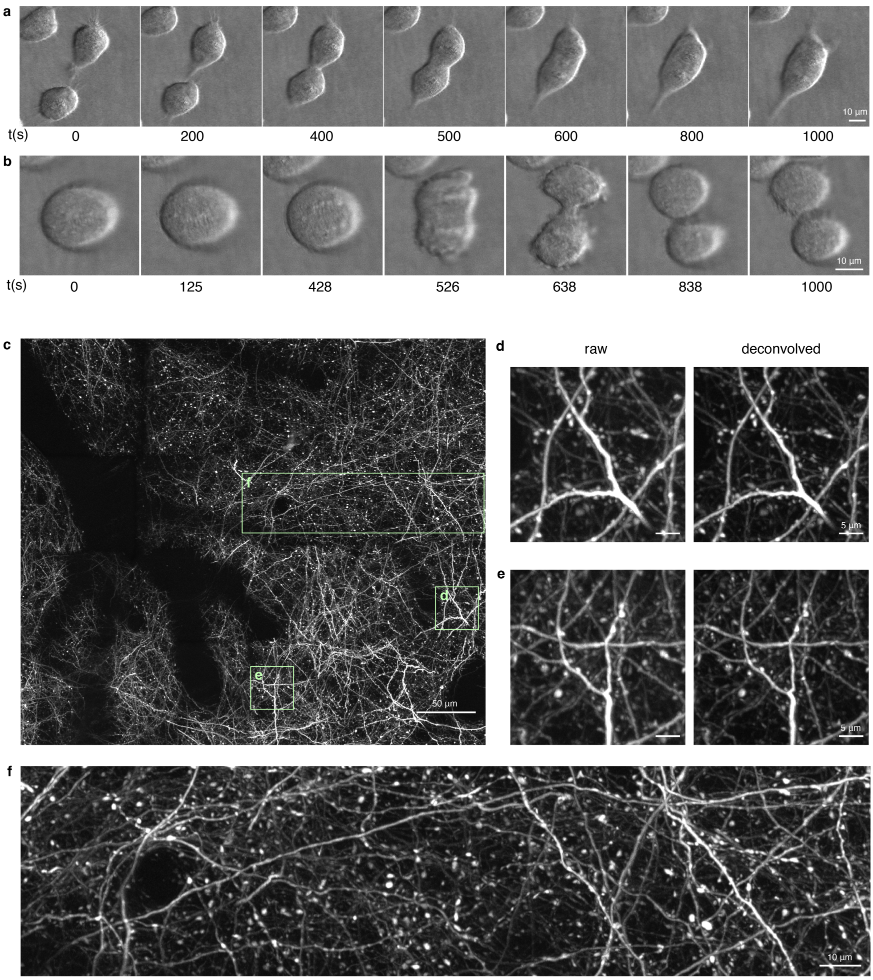


**Fig. S9**: Application of the processing pipeline to other imaging modalities. **a**, Time-lapse of a cropped region from the large field of view oblique illumination "phase" imaging of HeLa cells in Movie S1 showing two cells merging. **b**, a different region showing a dividing cell. **c**, xy MIP of a two-photon adaptive optical image of dendrites and axons across a large stitched field of view in the cortex of a live mouse. **d-e**, comparison of raw and deconvolution images in smaller subregions. **f**, Example of an axon extending across 210 $\mu$m field of view.

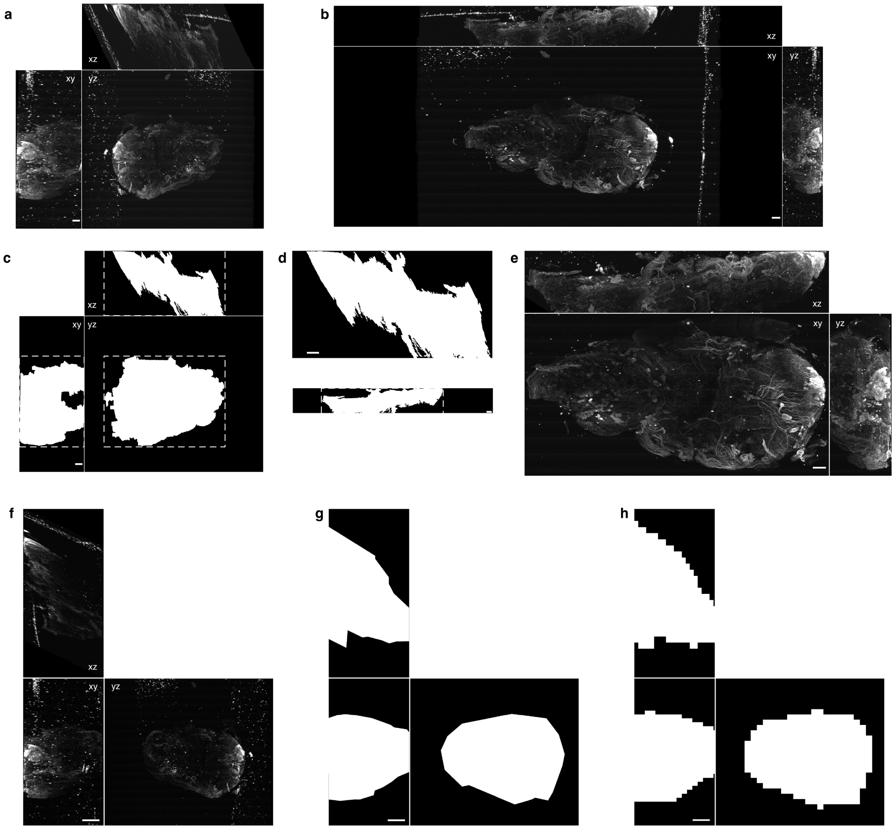


**Fig. S10**: MIP mask for efficient large-scale deskew/rotation and deconvolution. **a**, MIPs of the stitched data for the VNC data (Movie S4) in skewed space. **b**, deskewed and rotated results of the stitched data in **a** without MIP masking. **c**, segmented MIP masks from **a** that tightly bound the object for large-scale deskew and rotation. **d**, deskewed and rotated xz MIP to determine the tight bonding box for the output. **e**, the deskewed and rotated result with MIP masking. **f**, stitched data for the VNC data in skewed space. **g**, MIP masks generated by manual outline tracing. **h**, regions included for deconvolution after MIP masking based on the masks in **g**. Scale bars: 20 $\mu m$ for **a**-**e**, and 50 $\mu m$ for **f**-**h**.

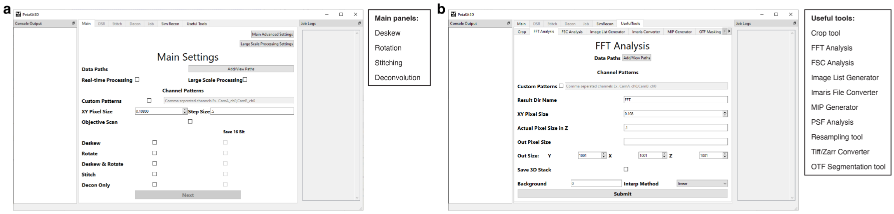


**Fig. S11**: Screenshots of GUI main panel and panel for useful tools. **a**, Screenshots of the GUI main panel with the list of processing steps. **b**, Screenshots of GUI panel for useful tools with the list of processing methods.

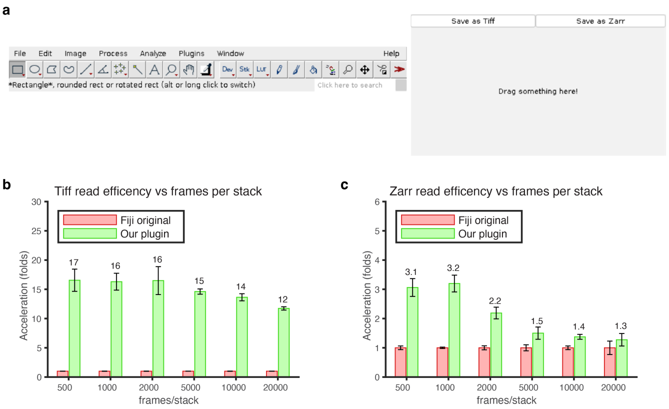


**Fig. S12**: Fast image reading using the Parallel Fiji Visualizer plugin. **a**, Screenshots of the Fiji main panel next to the drag-and-drop window of the plugin. **b**, Performance comparison a Tiff file (**b**) and a Zarr file (**c**) containing varying numbers of frames. The xy dimensions of the Tiff and Zarr images are $512 \times 1,800$. The Tiff files are compressed with the LZW algorithm, and the Zarr files are compressed with zstd algorithm with compression level 1. Benchmarks were performed on a computing node with dual Intel Xeon Gold 6146 CPUs, with three independent replicates for each test.

### Tables

| | TensorStore | | Cpp-Zarr |
|---|---|---|---|
| Input/output order | C-C | F-F | F-F |
| Reader (performance gain) | $1.00 \pm 0.04$ | $2.00 \pm 0.07$ | $2.25 \pm 0.07$ |
| Writer (performance gain) | $1.00 \pm 0.02$ | $0.0685 \pm 0.0012$ | $1.54 \pm 0.03$ |

**Table S1:** Comparison of reader and writer relative performance between TensorStore (in Python) and Cpp-Zarr (in MATLAB). Here different input and output data orders ("C": row-major layout, and "F": column-major layout) were benchmarked. For both reading and writing, "C-C" denotes operations from C to C layout, and "F-F" indicates F to F layout operations. Cpp-Zarr does not support "C-C" reading and writing operations. Performance gains are calculated by normalization over the "C-C" benchmarks for TensorStore. Benchmarks were performed on a $512 \times 1,800 \times 20,000$ uint16 image using a 24-core CPU node with dual Intel Xeon Gold 6146 CPUs, with each test repeated ten times.

| Dataset | Cells on coverslip | VNC | ExA-SPIM |
|---|---|---|---|
| #tiles | 4 | 1071 | 15 |
| Volume (TiB) | 0.035 | 27.8 | 107.54 |
| #nodes | 1 | 20 | 20 |
| Stitching-Spark (h) | $1.32 \pm 0.01$ | Not tested | Not tested |
| BigStitcher-Spark (h) | $0.109 \pm 0.003$ | $9.97 \pm 0.01$ | $20.1 \pm 0.2$ |
| New BigStitcher-Spark (h) [block scale] | $0.0234 \pm 0.0003$ [7,7,7] | $0.943 \pm 0.011$ [4,4,4] | $2.78 \pm 0.11$ [5,5,5] |
| Zarr-Stitcher (h) [batch size] | $0.0085 \pm 0.0003$ [1280,1280,512] | $0.333 \pm 0.001$ [1024,1024,256] | $1.40 \pm 0.00$ [768,768,768] |
| Fastest imaging time* (h) | 0.0094 | 12.6 | 17.6 |
| #node realtime processing* | 1 | 1 | 2 |

**Table S2:** Benchmarks of the three stitching methods. The table shows durations for the stitching (fusion) step, including both read and write times. The reported durations represent the actual time for the stitching step and do not account for total CPU hours (that can be calculated by multiplying with the total number of CPU cores employed). The benchmarks were performed on 24-core CPU nodes with dual Intel Xeon Gold 6146 CPUs. Stitching-Spark (version 402c45e) generated its registration information (not included in the presented processing time), while BigStitcher-Spark relied on the registration information from ZarrStitcher. We included two versions of BigStitcher: "BigStitcher-Spark" (version a942ffc) from late 2023, and "New BigStitcher-Spark" (version e20c64d), a substantially improved version from May 2024. The ExA-SPIM dataset originates from the work of Glaser et al. 2023. All benchmarks were run independently three times. The block sizes for BigStitcher and ZarrStitcher are $256 \times 256 \times 256$ and are $512 \times 512 \times 512$ for Stitching-Spark. To increase throughput, we fine-tuned the batch size (block scale in New BigStitcher-Spark and the batch size equals the block scale multiplied by the chunk size) for both New BigStitcher-Spark and ZarrStitcher. These optimal parameters for the benchmarks are listed under the corresponding running times. The benchmarks of Stitching-Spark on VNC and ExA-SPIM datasets could not be tested due to unsupported image readers. In the last two columns, we present the fastest data acquisition times, and the minimum number of computing nodes required for real-time stitching (assuming coordinate stitching without registration) for similarly sized data. *These estimates are based on an imaging frequency of 400 Hz (2.5 ms/frame) for cell and VNC data, and 6.1 Hz for ExA-SPIM (corresponding to the camera's max speed, VP-151MX-M6H00, Vieworks Co., LTD).

| #frame per stack | Imaging frequency (Hz) | Imaging time (s) | Read Tiff (s) | Deconvlution Deskew/ Rotate (s) | Write Zarr (s) | Total processing time (s) | #node real-time processing |
|---|---|---|---|---|---|---|---|
| 500 | 50 | 10 | 0.2 | 3.8 | 0.4 | 4.4 | 1 |
| 500 | 100 | 5.0 | 0.2 | 3.8 | 0.4 | 4.4 | 1 |
| 500 | 200 | 2.5 | 0.2 | 3.8 | 0.4 | 4.4 | 2 |
| 500 | 400 | 1.25 | 0.2 | 3.8 | 0.4 | 4.4 | 4 |
| 1,000 | 400 | 2.5 | 0.4 | 7.6 | 0.7 | 8.7 | 4 |

**Table S3:** Computing nodes (GPUs) required for real-time deconvolution and deskew/rotation for volumes with frame size $512 \times 1,800$ (xy) for different imaging conditions. The running times for the processing steps are taken from the figure panels Fig. S1a (Tiff read), S5c (deconvolution and deskew/rotation), and S1d (Zarr write) based on one CPU node with dual Intel Xeon Gold 6146 CPUs and one A100 GPU node.

| Figure/ Movie | Modality | $NA_{exc}$ | $\sigma_{NA}$ | $NA_{annulus}$ | $\Delta x_{sp}$ ($\mu m$) | time interval ($s$) | skewed image volume ($\mu m^3$) | # channel, # time, # tile |
|---|---|---|---|---|---|---|---|---|
| Fig. 2-3/ Fig. S1-S2/ Fig. S5i/ Fig. S12/ Movie S6 | Harmonic balanced hexagonal-rectangular lattice | 0.35 | 0.1 | 0.45/ 0.35 | 0.3 | N/A | 55×194×1020 | 1, 1, 1 |
| Fig. 4e/ Fig. S3a | Hexagonal multi-Bessel lattice | 0.43 | 0.08 | 0.47/ 0.40 | 0.23 | N/A | 38×52×100 | 1, 1, 1 |
| Fig. 4f | Swept sinc lattice | 0.32 | N/A | 0.40/ 0.20 | 0.27 | N/A | 38×52×135 | 1, 1, 1 |
| Fig. S3i | Harmonic balanced hexagonal-rectangular lattice | 0.50 | 0.15 | 0.60/ 0.40 | 0.27 | N/A | 38×52×100 | 1, 1, 1 |
| Fig. S3j | Swept Gaussian | 0.21 | 0.21 | 0.40/ 0.20 | 0.35 | N/A | 38×52×175 | 1, 1, 1 |
| Fig. 5c-e/ Fig. S9a-b/ Movie S1 | Oblique illumination (Phase) | N/A | N/A | N/A | N/A | 2.2 | 249×249 | 1, 1001, 20 |
| Fig. S6a | Widefield | N/A | N/A | N/A | 0.1 | N/A | 79×90×50 | 1, 1, 1 |
| Fig. S6c | Confocal | N/A | N/A | N/A | 0.1 | N/A | 86×86×50 | 1, 1, 1 |
| Fig. 5f-h/ Movies S2-S3 | Harmonic balanced hexagonal lattice | 0.40 | 0.10 | 0.50/ 0.35 | 0.26 | 89 | 55×194×1040 | 2, 301, 2 |
| Fig. S7 | ExA-SPIM | N/A | N/A | N/A | 1.0 | N/A | 10616× 7959× 8600 | 1, 1, 15 |
| Fig. S8a | Harmonic balanced hexagonal-rectangular lattice | 0.35 | 0.1 | 0.45/ 0.35 | 0.3 | N/A | 55×194×1020 | 2, 4, 4 |
| Fig. S9c-f/ Movie. S4 | Two photon | N/A | N/A | N/A | 1.0 | N/A | 111×111×50 | 1, 1, 32 |
| Fig. 6e-i/ Fig. S8b/ Fig. S10/ Movie. S5 | Harmonic balanced hexagonal lattice | 0.35 | 0.05 | 0.40/ 0.30 | 0.3 | N/A | 35×194×2472 | 2, 1, 1071 |

**Table S4:** Imaging conditions. The image data for Fig. 4e-f and Fig. S3a-c, g-j are from Liu et al. 2023. The image data for Fig. S7 comes from Glaser et al. 2023.

### Supplementary Movies (legends)

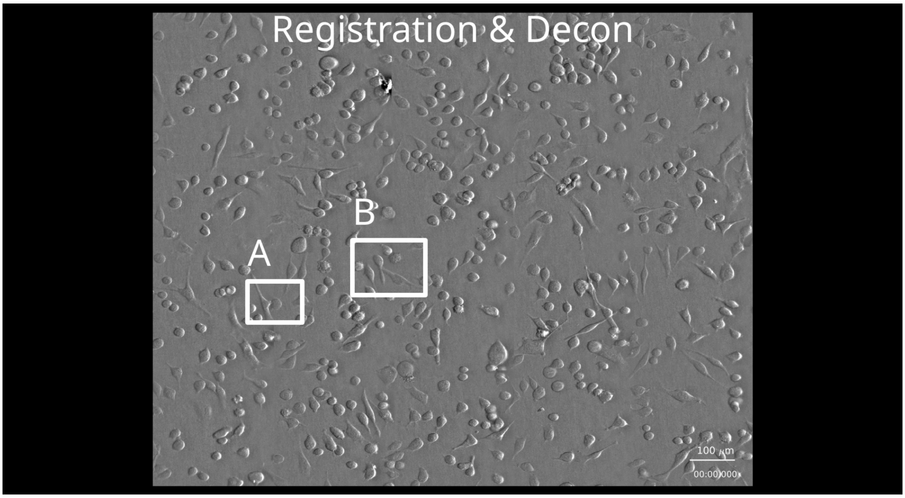


**Movie S1:** Large field of view oblique illumination "phase" contrast imaging of live HeLa cells over 1,001 time points at 2.2-second intervals. Here the deconvolution is compared with the situation without deconvolution. The stitching with or without flat-field correction and registration are also compared. The final processed data is 1,208 $\mu$m $\times$ 978 $\mu$m at each time point with a total size of 0.18 TiB.

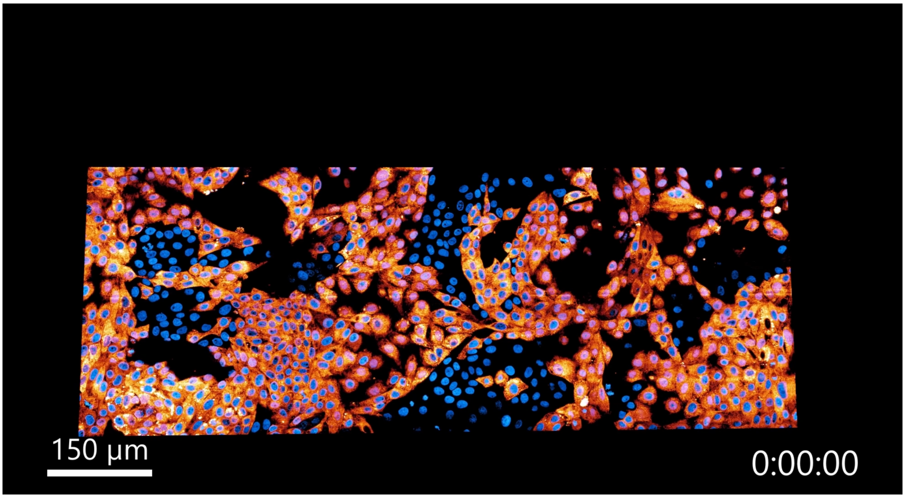


**Movie S2:** 3D large field of view live imaging of cultured LLC-PK1 cells by lattice light sheet microscopy for 301 time points at 89-second intervals. Orange: Connexin - mEmerald (ER), and blue: H2B - mCherry (nuclei). The final processed data for each channel is 1,015 $\mu$m $\times$ 378 $\mu$m $\times$ 15.2 $\mu$m at each time point with a total size of 5.1 TiB.

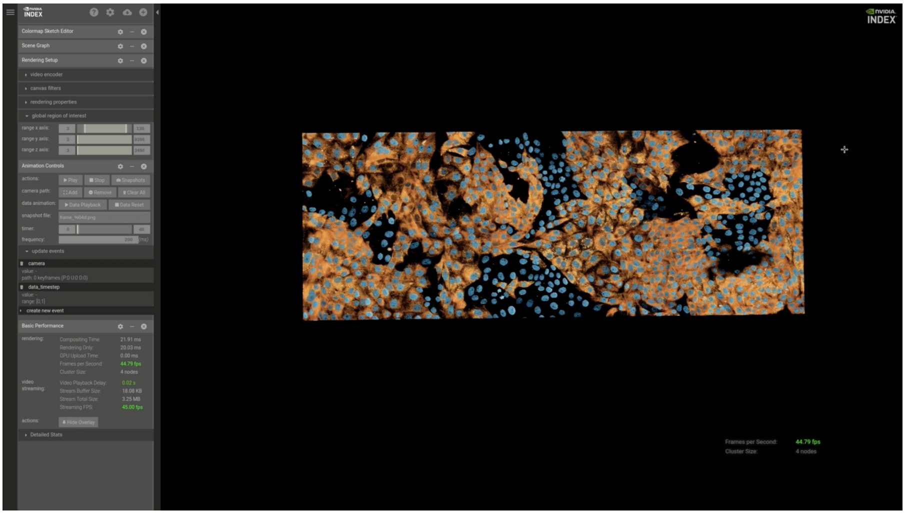


**Movie S3:** Screen recording of real-time 3D visualization of LLC-PK1 cells using NVIDIA-IndeX software in the uncompressed raw data format. This interactive visualization includes 40 time points (from 62 to 101 from Movie S2, with a total size of 0.67 TB). Orange: Connexin - Emerald (ER), and blue: H2B - mCherry (nuclei). The screen recording is sped up by 2.5 times.

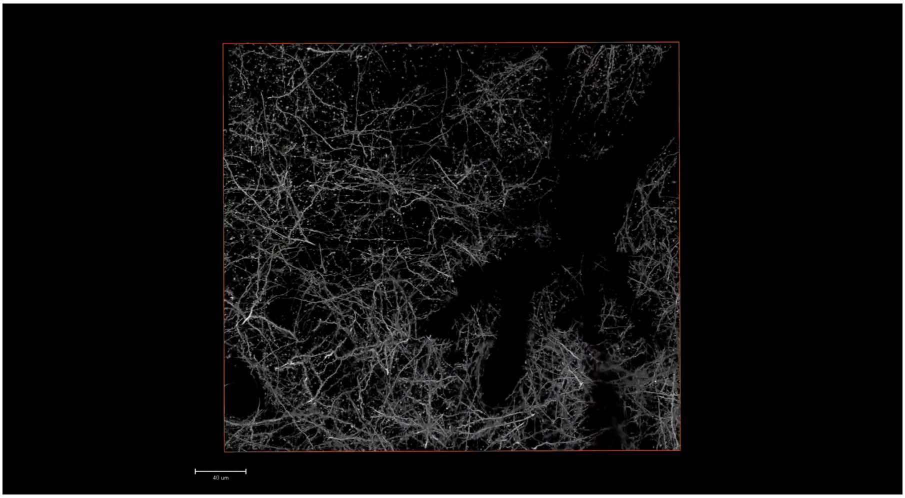


**Movie S4:** Movie through a large field of view live in the cortex of a live mouse, as seen by adaptive-optical two-photon microscopy. The final processed data is 380 $\mu$m $\times$ 333 $\mu$m $\times$ 10.9 $\mu$m with a total size of 2.0 GiB.

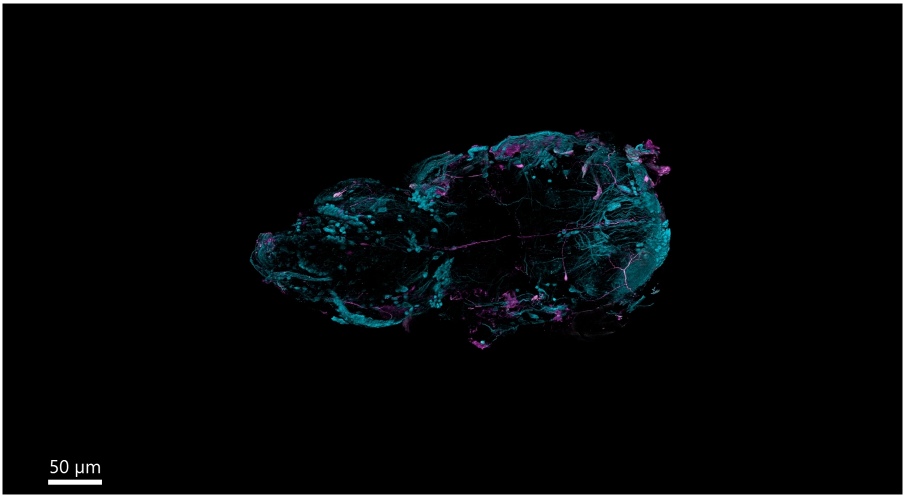


**Movie S5:** Movie of a whole fly VNC imaged at 8× expansion. Cyan: VGlut$^{MI04979}$-LexA::QFAD, and purple: MN-GAL4. The final processed data for each channel is 3,856 $\mu$m × 1,750 $\mu$m × 902 $\mu$m with a total size of 9.9 TiB.

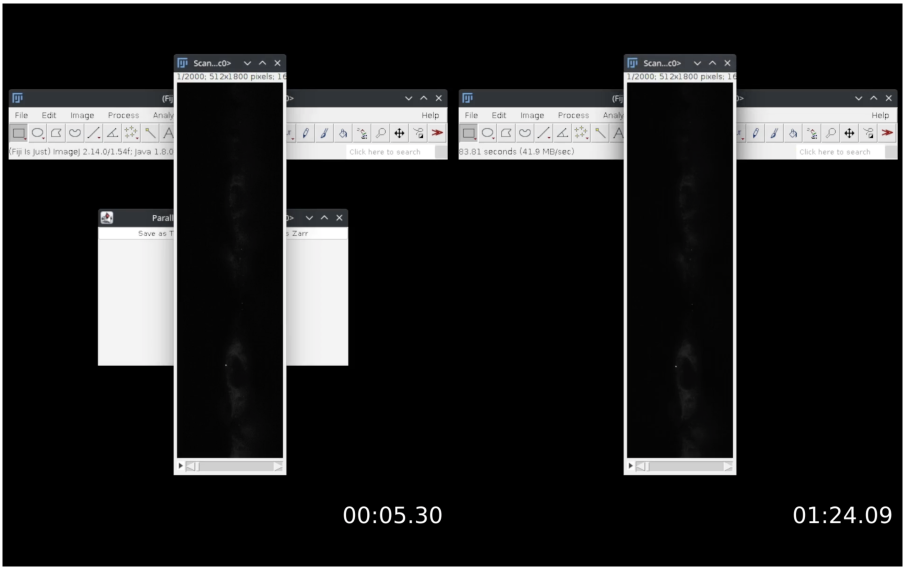


**Movie S6:** Screen recording of opening the same LZW compressed Tiff file using the Parallel Fiji Visualizer plugin (left) and the native Fiji reader (right). Tests were performed on $512 \times 1,800 \times 2,000$ uint16 image (2.0 GiB compressed and 3.4 GiB uncompressed) using a 24-core CPU node with dual Intel Xeon Gold 6146 CPUs.
